# The Metabarcoding Analysis Pipeline (MAP): Simple, accurate, and flexible metabarcoding

**DOI:** 10.64898/2026.07.22.740107

**Authors:** Sean WJ Prosser, Nicholas W Bard, Ken A Thompson, Robin M Floyd, Sameer Padhye, Emine Ozsahin, Saeideh Jafarpour, Paul DN Hebert

## Abstract

Current metabarcoding pipelines are inflexible with respect to study design and are poorly suited to long-read sequence data. To address these limitations, we developed MAP, the Metabarcoding Analysis Pipeline, which is a sequence-to-answer workflow supporting the analysis of amplicons from highly multiplexed and replicated study designs. Although MAP can analyze amplicons of any length from any genetic marker, it includes several features tailored to long-read COI metabarcoding. MAP installs from a Docker container and requires only sequence data, a parameters file, and a reference library. It produces intuitive reports, enabling users to evaluate their data immediately after analysis. We validate MAP by showing that it generates biodiversity estimates that correspond closely to a ground-truth dataset of single-specimen DNA barcode data and by demonstrating that it outperforms alternative platforms for COI metabarcoding. MAP is free, open-source, and available from: https://github.com/cbg-innov/MAP.

## INTRODUCTION

Biodiversity monitoring is critical for quantifying changes in the abundance and distribution of species (Janzen *et al*., 2020). For large-bodied or vocal taxa, cost-effective monitoring can be achieved through camera trapping (Ahumada *et al*., 2020) and soundscape analysis (Müller *et al*., 2023). However, these species represent a tiny fraction of all life (Stork, 2018; Colwell *et al*., 2026); for the remaining taxa, the silent and small, alternative approaches are needed to document their presence and abundance (Müller *et al*., 2023). The past 20 years have seen increasing reliance on DNA-based identification tools, which match a small region of an organism’s genome to a reference database to generate an identification (Hebert *et al*., 2003). Since these approaches—termed ‘DNA barcoding’ when applied to single specimens and ‘metabarcoding’ when applied to bulk samples containing multiple specimens—are critical for monitoring most species, optimizing protocols for accuracy and cost-effectiveness is essential.

DNA barcoding and metabarcoding rely on sequencing DNA from defined genomic regions, and matching those sequences against a reference library of sequences derived from specimens with known taxonomy (Blagoev *et al*., 2016; Fontes *et al*., 2021). DNA barcoding reliably associates full-length barcode sequences with specimens because one only expects a single sequence and those that do not match the expected taxonomy can be filtered out or marked as non-target sequences (Mioduchowska *et al*., 2018). DNA barcodes are centrally archived and curated on the Barcode of Life Data System (BOLD) (Ratnasingham and Hebert, 2007; Ratnasingham *et al*., 2024). While DNA barcoding originally relied on Sanger sequencing, advances in long-read technology via next-generation sequencing (NGS) platforms have spurred a shift to long-read NGS for producing DNA barcodes (Hebert *et al*., 2018, 2024).

Metabarcoding is used for biodiversity assessment because it is less expensive than barcoding, but it presents methodological challenges. Sequences recovered from a metabarcoding sample typically derive from many species and must be computationally clustered (Taberlet *et al*., 2012). Metabarcoding also raises the risk of technical artefacts. For example, chimeric sequences—formed when taxonomically distinct amplicons combine during PCR—must be identified and removed (Edgar *et al*., 2011). Tag jumps also complicate the interpretation of results (Schnell *et al*., 2015). Contamination filters must be fine-tuned because low-frequency OTUs can represent true signal from rare or low-biomass specimens—particularly those with hard exoskeletons (Clare *et al*., 2016)—rather than contamination or sequencing error. One important category of artefacts, nuclear mitochondrial DNA segments (NUMTs), are much less prevalent when PCR targets long amplicons (Hebert *et al*., 2023). However, metabarcoding has generally relied on short-read sequencers (Furneaux *et al*., 2025), largely because the sole long-read platform (Sequel II) delivered too few reads to be cost-effective. Because ONT’s PromethION flow cell generates up to 200 million reads, it has removed this constraint, but there has, until now, been no software designed to analyze these data.

This paper introduces MAP, the Metabarcoding Analysis Pipeline, which supports the analysis of metabarcoding data generated by long– or short-read sequencers. MAP is the core metabarcoding informatics pipeline used at the Centre for Biodiversity Genomics. It processes reads from a metabarcoding sequencing run, clusters them into OTUs, and assigns taxonomy to each OTU by comparison with a reference library (e.g., BOLDistilled libraries) (Prosser *et al*., 2025). MAP is sequence-length agnostic, can analyze reads from any genetic marker, support any number of samples, and accommodate any level of per-sample replication. The balance of this publication introduces MAP and validates its performance by analyzing a dataset generated by barcoding every specimen present in ten samples that were also metabarcoded via both long– and short-read sequencers. Using these results, MAP’s performance is benchmarked by comparing its output with those generated by alternative platforms.

## METHODS

### MAP Software

MAP facilitates the analysis of metabarcoding data because of its flexibility with respect to markers, primers, unique molecular identifiers (UMIs), and sequencing platforms. This paper summarizes key methodological components of MAP v1.0.1 while a complete description, which will be updated regularly to reflect software enhancements, is available online at: https://github.com/cbg-innov/MAP.

MAP requires four inputs to run: (1) a ‘parameters’ worksheet in *.xlsx format (built in Microsoft Excel or equivalent) which assembles sample metadata and run-wide and sample-specific parameters; (2) raw sequence data (*.fq.gz); (3) a reference library or libraries (*.fasta) for the barcode(s) used; and (4) a compose.yaml file for the working directory, which is provided. A second reference library is included with MAP that is used exclusively for sequence correction when the target sequence is ≥ 500 bp COI-5P. The parameters file is flexible, allowing the user to implement study designs involving any number of samples, multiple markers and/or primers, and varying numbers of technical replicates, making it possible to implement best practices for metabarcoding study design (Zinger *et al*., 2019). See Supplementary Text for more details on Sequence Correction and Reference Panel creation.

We recommend installing MAP using Docker (available at https://www.docker.com/) with the image file provided in GitHub Packages at: https://github.com/cbg-innov/MAP/pkgs/container/map. Running a container from the Docker image file generates MAP’s working environment for successful command execution and installs all software required to complete each stage of the workflow. These programs are version-controlled to prevent incompatibilities but can be customized by the user if desired. Running the Docker container assembles the correct directory structure and downloads and unpacks a reference library, which are both required to run MAP. Its COI workflows have been tested with BOLDistilled libraries (Prosser *et al*., 2025) which are downloadable, comprehensive COI barcode libraries that are 50× faster to query than the full BOLD library yet generate near identical results. Alternatively, users can provide their own custom reference libraries for other markers and/or for COI. Reference libraries must be provided as SINTAX-formatted .fasta files (Rognes *et al*., 2016).

First-time users should verify successful installation by performing a complete analysis of the test data set upon startup, which can be achieved by running the container without supplying new data or additional arguments on the command line. After running the container for the first time, users can work inside it, edit or replace the pre-existing parameters worksheet (*.xlsx), supply their own raw data (*.fq.gz), and provide and/or replace the reference library (*.fasta) files. After successful initialization, users may exit and return to the Docker container using its stop/start commands. The default reference library is stored in a Volume and saved locally for future sessions. Experienced users can run MAP directly using custom files by adding them to the /Docker directory and specifying filenames as command-line flags.

Once installed, MAP is called via the command line. It automatically detects the number of CPU cores available to the user (by default, using all but two cores) and leverages parallel processing when possible. The number of cores used by MAP can also be adjusted by the user. Processing time varies with the number of reads, sequence diversity, sample/replicate quantity, and hardware.

A full visual walkthrough of MAP is presented in Fig. 1 with some specific steps varying according to the data source. Briefly, MAP merges R1 and R2 input sequence files if Illumina data are supplied using `pear` (Zhang *et al*., 2014)., Next, MAP uses `chopper` (De Coster and Rademakers, 2023) to remove aberrantly long or short ONT reads, or `vsearch ––fastq_filter`, to remove high error Illumina merged reads. Next, MAP and demultiplexes, trims primers, and size-selects sequences in parallel (Tange, 2025) using cutadapt (Martin, 2011); formatting and reverse-complementation assistance is provided by seqtk (Li, 2025) and seqkit (Shen *et al*., 2016). By default, MAP employs a 12.5% error rate for UMIs acquired with nanopore sequencers and a 0% error with Illumina; both can be changed by the user. A 20% error rate in primer sequences is permitted; since primers are typically very distinct, OTUs assigned to the incorrect marker are unlikely and are easily recognized should they occur. Both termini are searched for the forward and reverse primer sequences (unless using paired-end reads); and backwards reads are reverse complemented when present.

**Fig. 1.**
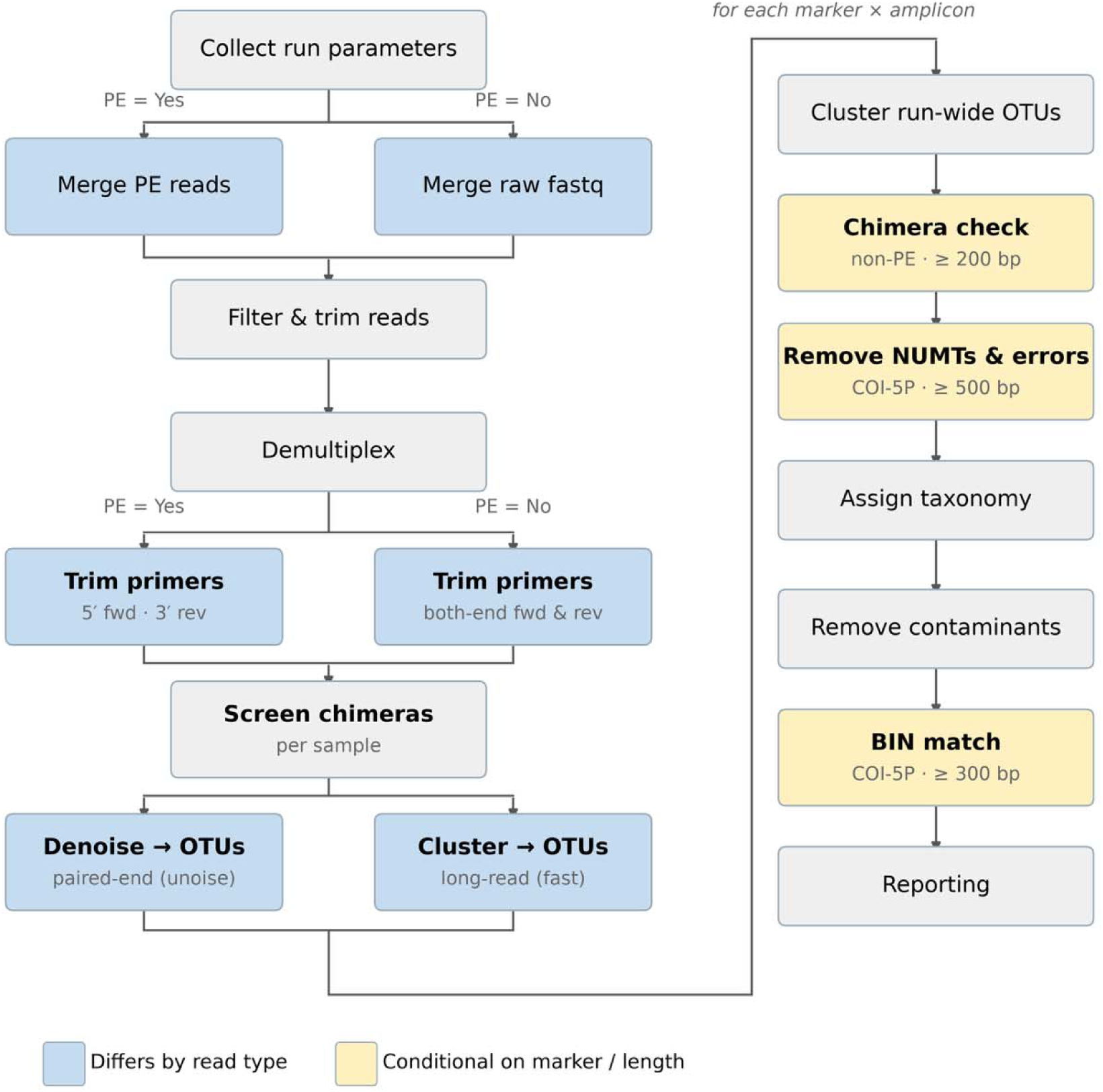
Workflow diagram for MAP. Some steps are invoked only when analyzing specific data types or markers, as indicated.

Following these steps, reads are clustered into OTUs using vsearch –– cluster_fast, and only those that meet the user-defined minimum read-depth threshold are retained (we suggest 3 for nanopore and 2 otherwise). Multiple forward or reverse primers (i.e., primer cocktails) are supported for each marker in a particular run. Put differently, MAP can process a run containing any number of primers for any number of markers. The user has the option to employ the same UMI adapters in different combinations to mark different samples by checking for linked primers with cutadapt.

Amplicons are dereplicated, and, if using Illumina paired-end read data, those ≥200 bp are screened for chimeras using vsearch ––uchime_denovo. A chimera check for long-read sequence data is implemented following run-wide clustering (Hakimzadeh *et al*., 2025; Overgaard *et al*., 2024). The filtered amplicon sequences are next clustered into sample-specific OTUs based on sequence similarity (*i.e.,* “primary clustering”). The user may elect to retain the component reads for the OTUs formed at this stage.

While MAP supports the analysis of any genetic marker, certain features are only available for amplicons of a particular length and/or for COI. For example, if using the standard animal barcode region (COI-5P), the OTU consensus sequences are automatically trimmed to the precise COI barcode region using an alignment-based approach after OTU clustering and filtering. To qualify, the target amplicon must be COI-5P and ≥500 bp. Samples that do not qualify—which includes most data generated by short-read sequencers—bypass this step. Consensus sequences generated via nanopore sequencers primarily contain errors in homopolymer regions with ≥4 identical adjacent bases. MAP’s error correction module for COI-5P aims to identify sequencing errors when an incorrect number of bases was assigned in homopolymer tracts during basecalling or consensus assembly. Briefly, MAP aligns sequences to a customized standard reference panel (see Supplementary Methods – Reference Panel Creation), identifies 1-bp or 2-bp errors within homopolymers, and either fills those gaps with ambiguous (‘N’) bases, or deletes extra bases, so they align in the proper reading frame. At this stage, long-read sequence data are screened for chimeras using vsearch uchime_denovo.

Following these COI-specific steps, clustered OTUs are pooled across all samples to generate run-wide OTUs. These run-wide OTUs are assigned taxonomy probabilistically using vsearch ––sintax (Edgar, 2016) and potential contaminant OTUs from each sample are identified and removed as follows (‘the contaminant threshold test’). First, read-depth ratios are computed for each OTU and those exhibiting counts consistent with cross-contamination are flagged. To distinguish between contamination and genuine low-level detections, MAP determines a background noise threshold by inspecting the abundance of each OTU in negative control wells compared to wells with samples. First, OTUs observed with higher read counts in negative controls than in samples are removed from all samples. For the remaining OTUs detected in negative controls, a basic minimum read depth threshold is determined by multiplying the proportion of control to sample read counts by a customizable parameter value (α-quantile, default = 0.9). Using this noise threshold, a minimum read depth is calculated independently for each OTU, and detections supported by fewer reads are tagged as possible contaminants but not yet discarded.

MAP considers the distribution of OTUs among technical replicates (if used) to aid its denoising decisions, a feature which helps to determine if OTUs with few reads are genuine or artefacts. An additional ‘replicate test’ is applied to potentially retain rare and/or low abundance OTUs that were tagged as possible contaminants. The replicate test consists of two components, for which each OTU must meet all criteria within a sample to pass. In the first component, the OTU must occur in a preset minimum proportion of replicates (min_rep_prop default value = 0.65; with eight replicates presence in > 5). Because contamination is likely to occur at different levels among sample replicates, the next component tests for the consistency of read depths among replicates. The sequence depth distribution is assessed for designated OTUs present in at least three replicates, and the criteria is met if a preset minimum proportion of sample replicates occur at similar sequence depth values. The similarity in sequence depth is determined by calculating the kernel density of the sequence depth distribution using the Sheather-Jones algorithm (Sheather and Jones, 1991). The minimum and maximum values for the sequence depth “hot zone” are determined by applying *n* Jenks natural breaks (based on replicates present per OTU) to the entire distribution. Samples that do not have a minimum proportion of their replicates in which the OTU occurs within the same Jenks natural break (high_dens_prop default = 0.65) do not meet the criteria for this component and fail the replicate check. If both the ‘contaminant threshold’ and ‘replicate’ tests fail, the OTU for the sample is discarded. Users can tune these parameter values to their preferences and defaults will be refined as further datasets become available for exploration.

For COI-5P amplicons ≥ 300 bp, MAP performs a Barcode Index Number (BIN) matching analysis. BINs (Ratnasingham and Hebert, 2013) are sequence clusters that delimit species and are highly congruent with morphological taxonomy in many animal taxa (Ortiz *et al*., 2017; Prieto *et al*., 2021). OTUs are matched to BINs using vsearch ––usearch_global. For this step, the user must have included BIN designations in their original SINTAX-formatted reference library. MAP will automatically create a properly formatted library for vsearch ––usearch_global if the user does not have one. Because the initial BIN clustering threshold is 2.3% (Ratnasingham and Hebert, 2013), we adopted this value as the maximum distance for query sequences to be considered a BIN match; otherwise, they are deemed to lack a match. For OTUs that do not match a known BIN, the nearest neighbour BIN information is returned. For all BIN-matched OTUs, MAP returns a distance value from the corresponding BIN sequence.

Finally, results are provided on a per-sample and per-replicate basis in a single Excel (.xlsx) file optimized for readability. A summary report (.html), generated using quarto (Allaire and Dervieux, 2025), provides an overview of run statistics including read counts, demultiplexing success, OTU counts, and taxonomic composition. The following stand-alone software packages were also used by MAP: Bioconductor (Huber *et al*., 2015), biopython (Cock *et al*., 2009), biostrings (Pagès *et al*., 2025), iNEXT (Hsieh *et al*., 2016). The following R (R Core Team, 2024) packages were also invoked: `data.table` (Barrett *et al*., 2025), `DT` (Xie *et al*., 2025), `htmltools` (Cheng *et al*., 2024), `kableExtra` (Zhu, 2024), leaflet (Cheng *et al*., 2025), `paletteer` (Hvitfeldt, 2021), `openxlsx` (Schauberger and Walker, 2025), `plotly` (Sievert, 2020), `shiny` (Chang *et al*., 2025), `tidyverse` (Wickham *et al*., 2019), `treemapify` (Wilkins, 2023).

### Validation: Muogamarra Case Study

We validated the effectiveness of MAP via the analysis of an original dataset. Ten Malaise trap samples were collected near the Muogamarra Nature Reserve, New South Wales, Australia (−33.59920, 151.16429) in December 2024. Samples were brought to the Centre for Biodiversity Genomics for analysis under permit PWS2024-AU-002999.

All ten samples were initially metabarcoded and their component specimens were then individually barcoded. Briefly (see Supplementary Methods for full details), DNA was extracted from eight technical replicates per sample, and the 658 bp COI barcode region was amplified using symmetrical indexed primers. This resulted in 80 amplicon pools while eight positive and eight negative controls filled the remaining wells in a 96-well plate. Negative controls were used for denoising as described above, while positive controls were used to confirm positive detections via a synthetic control and then ignored. Amplicons were sequenced on an ONT PromethION 2 Solo sequencer. The run generated 179,806,945 base-called reads and was analyzed using MAP v1.0.1. (see Data Accessibility for access to input files). To compare results with short-read data, the same extracts were also amplified for a ∼418-bp COI amplicon using the BF3/BR2 primers (Elbrecht and Leese, 2017; Elbrecht *et al*., 2019) on an Illumina NextSeq P2 flow cell that was shared with other projects. The run generated 67,324,112 reads for the focal samples, which were processed via MAP v1.0.1. A follow-up study will explore the similarities between the Nanopore and Illumina datasets in greater depth.

After metabarcoding, every specimen from each bulk sample was individually barcoded. We photographed each specimen before extracting DNA from it and then conducted PCR using the same primers employed for ONT metabarcoding. We sequenced all amplicons on a MinION Mk-1d sequencer and conducted three additional runs with specimens that initially failed to return a sequence to maximize barcode recovery. Barcode results are available on BOLD as a public dataset DS-PHAUS24 (dx.doi.org/10.5883/DS-PHAUS24). We assigned BINs and taxonomy to the barcode results via MAP’s BIN matching module using the April 2026 COI BOLDistilled library (Prosser *et al*., 2025). Barcode data were analyzed using BIP (Prosser et al., 2026).

BIP outputs both barcodes for target specimens and non-target sequences (NTS; e.g., parasitoids, stomach contents). For most barcode data in the main text, we present analyses that compare metabarcode data to both target and NTS together. We present an analysis of only target sequences in the supplementary materials. This approach is justified because NTS are expected to be present in metabarcode data, so their detection represents a true positive. Excluding them from consideration would penalize accuracy when evaluating the performance of MAP to other pipelines. We consider the single-specimen barcode data as the ‘ground truth’. We retained only unflagged records with full-length barcodes, ≥ 97.7% match to a BIN, and <1% ambiguous bases. For the barcode dataset and all metabarcode datasets, we filtered data to only include the phylum Arthropoda and removed sequences that could not be assigned to a class.

We evaluated model performance by computing biodiversity metrics routinely employed to quantify α*-,* β-, and γ-diversity. For α*-*diversity, we computed BIN richness of each sample and then compared it to the ground truth estimates. We then computed the Concordance Correlation Coefficient (CCC) (Lawrence and Lin, 1989) which evaluates the degree to which the two vectors have a slope of 1 and an intercept of 0. We also report the Spearman rank-order correlation of BIN richness between barcode and metabarcode datasets. For β-diversity, we computed the Bray-Curtis distance matrix among samples based on the presence-absence of BINs. We then used a Mantel test to evaluate the correlation between the barcode and metabarcode matrices (Legendre *et al*., 2008). We also report the mean Bray-Curtis distance (±SD) between the barcode and metabarcode data. Visualization proceeded via multidimensional scaling (MDS). Finally, for γ-diversity, we output the F1 score across the entire dataset (Powers, 2020) which evaluates a model’s ability to correctly identify true taxa (recall) while minimizing incorrect detections (precision) when compared against the barcode dataset. We also report the percentage of target (i.e., non-NTS) BINs that were present in the metabarcode dataset.

We acknowledge that the present validation test is based on a single dataset. However, obtaining this dataset was expensive because it involved barcoding more than 11,000 specimens with multiple attempts to recover failures, and sequencing on both Illumina and ONT. Thus, while this is a limitation, it is one imposed by the cost in implementing a rigorous validation with large samples.

Metabarcoding datasets are ordinarily filtered before downstream analysis to remove low frequency sequences that derive from contamination or PCR/sequencing errors (Zinger *et al*., 2021). We present all analyses under three filtering regimes. First, we simply present the unfiltered output. Second, we use a permissive proportional filter that retains any OTUs with ≥0.0001% of all reads from that sample. Last, we use a more stringent proportional filter that retains any OTUs with ≥ 0.001% of all reads from that sample.

### Benchmarking

We compared MAP’s performance to leading alternative pipelines for the analysis of COI metabarcoding data. Although there are a ‘pile of pipelines’ (Hakimzadeh *et al*., 2024) for metabarcoding, MAP was developed to address gaps in these pipelines. Specifically, no existing pipeline accepts long-read data, is tailored for COI barcoding, and supports the full analytical path from raw sequence data to an OTU table with BIN assignments (Table 1). We specifically compared MAP to the two other platforms that return BIN assignments using reference libraries: mBRAVE (mBRAVE.net) and spcfy.io. We also compared MAP to MetaWorks (Porter and Hajibabaei, 2022) and QIIME2 (Bolyen *et al*., 2019), and used BINs returned from MAP’s BIN-matching module for these analyses. We did not use DADA2 (Callahan *et al*., 2016) because it returns ASVs, not OTUs. We attempted to use PRONAME (Dubois *et al*., 2024) to analyze the ONT data, but its storage and memory requirements precluded its application to PromethION-scale data. mBRAVE and spcfy.io are hosted online and analysis is completed on their servers whereas the others are precompiled and run on the user’s computational resources. mBRAVE and spcfy.io have not been peer reviewed while the other pipelines are published. While mBRAVE can support sequences up to 1500 bp and could potentially analyze our ONT data, the demultiplexed files for our PromethION run were too large to upload. See the Supplementary Text for specific details about our use of these platforms. BIN assignments were made using MAP’s BIN matching module for all pipelines except mBRAVE, which does not provide sequences. spcfy.io does provide BINs, but we re-assigned BINs to match the most current reference library (this improved its performance).

**Table 1.**
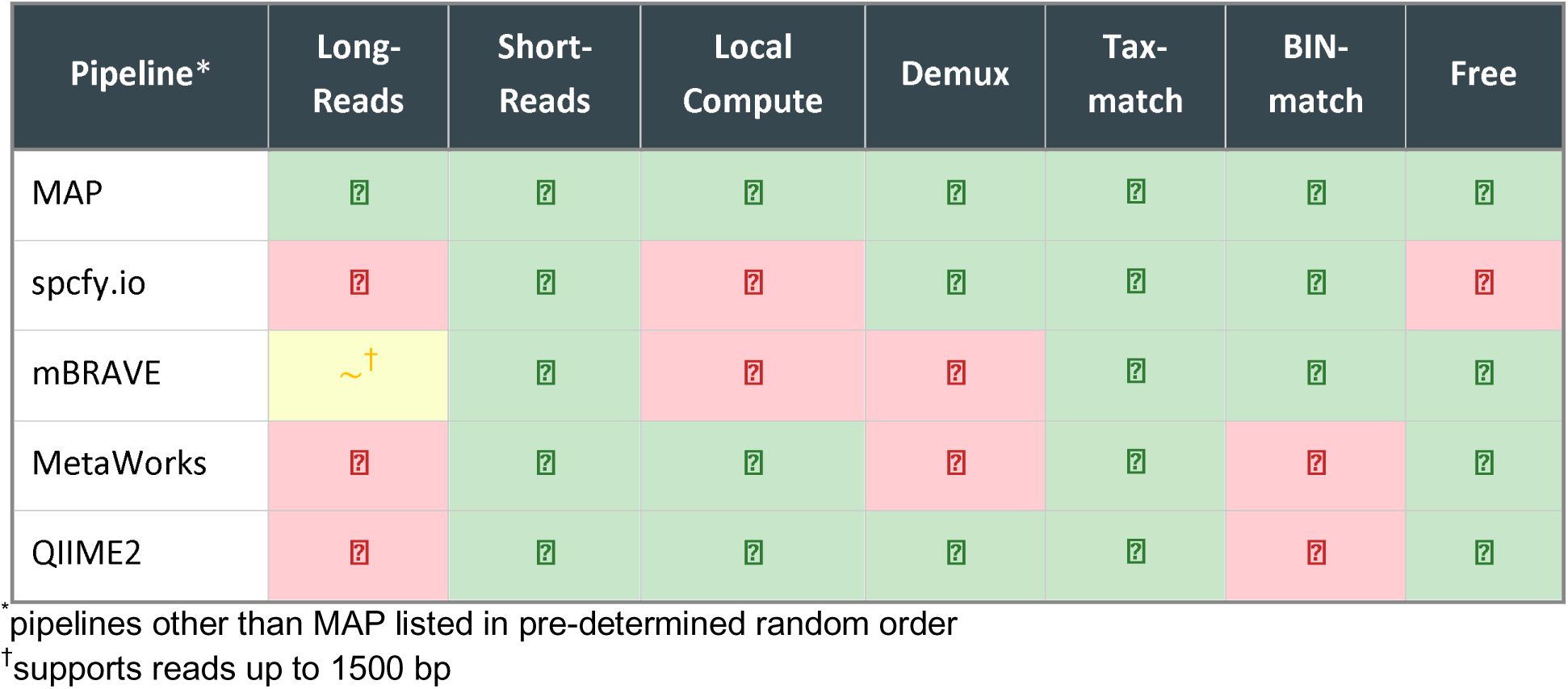
Comparison of MAP’s features to those of other metabarcoding analysis pipelines (as of July 2026).

### Data analysis

Data analysis used custom scripts which are available online (see Data & Code Accessibility). Data import, visualization, and/or wrangling was facilitated by the following R packages: BOLDconnectR (Padhye *et al*., 2025), readxl (Wickham and Bryan, 2019), and tidyverse (Wickham *et al*., 2019). Computation of β-diversity metrics used functions in vegan (Dixon, 2003). Data visualization invoked eulerr (Larsson and Gustafsson, 2018), ggalluvial (Brunson and Read, 2023), patchwork (Pedersen, 2019), and scales (Wickham *et al*., 2025).

## RESULTS

### Barcoding results

The ten bulk samples contained 11,509 specimens, and we obtained high-quality (BIN-eligible) barcode sequences from 10,903—a success rate of 94.7%. We found a total of 1,700 BINs among these specimens. Of these, 1,104 were detected only as target (parent) records, 57 only as non-target sequences (NTS), and 539 were detected as both. Taxonomically, 1,557 BINs (91.6%) were insects, 87 (5.1%) were arachnids, 55 (3.2%) were collembolans, and one was a millipede.

### Validation of MAP with ONT Data

Biodiversity metrics produced by MAP when analyzing ONT data are presented in Table 2 for the three filter strategies (Figures 2 & S1). Regardless of filter strategy, α-diversity estimates from MAP were highly correlated with those from the barcode dataset. For β-diversity, MAP’s distance matrices were an accurate representation of those determined via single-specimen sequencing, though estimates were greatly improved by filtering. γ-diversity estimates were also accurate: most target BINs were detected and MAP yielded consistently high F1-scores.

**Fig. 2.**
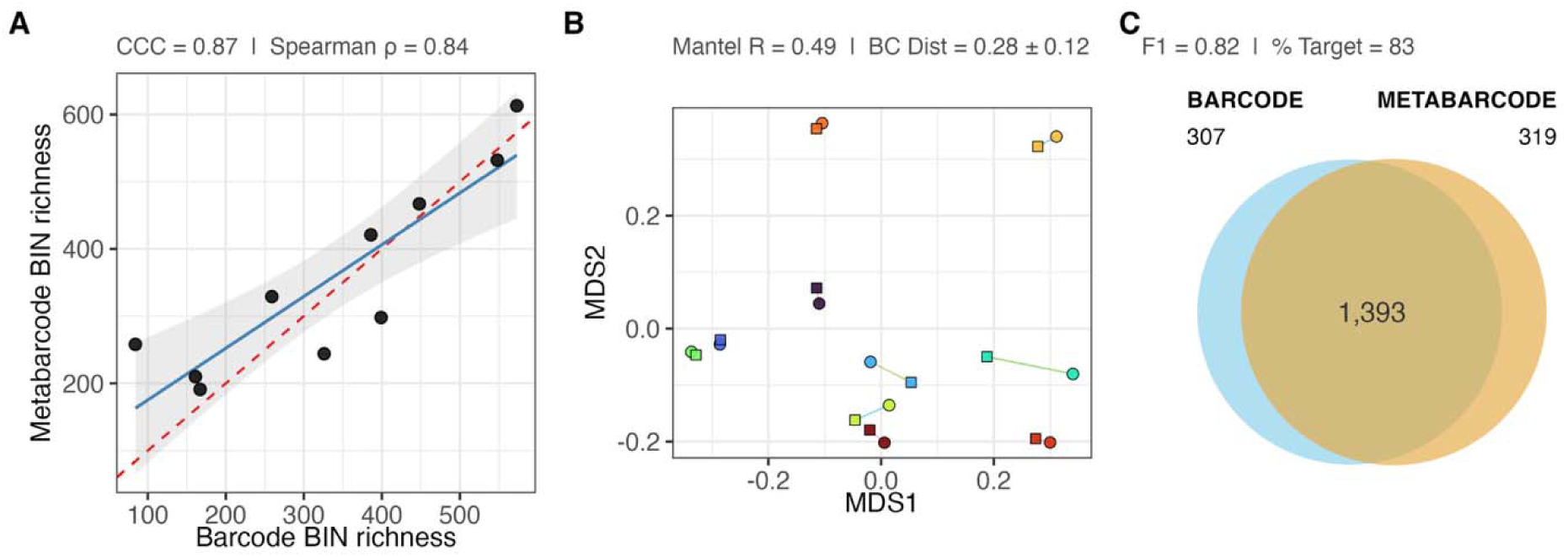
Visualization of biodiversity metrics for MAP analysis of unfiltered ONT data based upon the comparison of barcode and metabarcode datasets. (A) α-diversity metrics evaluate the degree to which per-sample BIN richness estimates from barcoding and metabarcoding correspond. CCC evaluates the correspondence to 1:1 with an intercept through the origin; Spearman’s ρ evaluates the correlation of rank-orders. (B) β-diversity visualized as an MDS plot; circles are barcode and squares are metabarcode results. Mantel’s *R* evaluates the correlation between distance matrices while Bray-Curtis distance evaluates the average dissimilarity. (C) γ-diversity is estimated across the entire dataset; in the Venn diagram numbers indicate the BINs detected in both datasets vs. those unique to either the barcode or metabarcode dataset. F1 score evaluates the balance of precision (avoiding false positives) and recall (avoiding false negatives). % Target indicates the percentage of target barcodes in the metabarcode dataset. Similar figures for other pipelines and filtering strategies and datasets are provided in the Supplementary Materials. Perfect matches between the barcode and metabarcode datasets yield a value of ‘1’ for all metrics except for BC distance, where a perfect match yields a value of ‘0’.

**Table 2.** Comparison of MAP’s performance to other pipelines using three data filtration strategies; darker green indicates better performance within each metric and filter strategy.

| Filter Strategy | Software | Dataset | Min. Reads | Min. Reps | # Reads in OTUs | CCC ($\alpha$) | Spearman $\rho$ ($\alpha$) | Mantel $R$ ($\beta$) | BC Dist ± SD ($\beta$) | % Target ($\gamma$) | F1-score ($\gamma$) |
| --- | --- | --- | --- | --- | --- | --- | --- | --- | --- | --- | --- |
| Unfiltered | MAP | Nanopore | 3 | 1 | 118.1M | 0.87 | 0.84 | 0.49 | 0.28 ± 0.12 | 83 | 0.82 |
| | MAP | Illumina | 2 | 1 | 43.8M | 0.75 | 0.82 | 0.32 | 0.38 ± 0.10 | 85 | 0.73 |
| | spcfy.io | Illumina | 1 | 1 | 37.5M | 0.29 | 0.71 | 0.23 | 0.57 ± 0.10 | 71 | 0.59 |
| | mBRAVE | Illumina | 1 | 1 | 56.7M | 0.059 | 0.85 | 0.041 | 0.60 ± 0.14 | 89 | 0.69 |
| | MetaWorks | Illumina | 3 | 1 | 59.8M | 0.16 | 0.81 | 0.21 | 0.62 ± 0.10 | 74 | 0.51 |
| | QIIME2 | Illumina | 2 | 1 | 44.2M | 0.24 | 0.85 | 0.23 | 0.59 ± 0.096 | 74 | 0.53 |
| Permissive filter ($\geq 0.0001\%$ of sample reads) | MAP | Nanopore | $\geq 4$ | 1 | 117.9M | 0.87 | 0.89 | 0.53 | 0.27 ± 0.11 | 80 | 0.81 |
| | MAP | Illumina | $\geq 3$ | 1 | 43.8M | 0.81 | 0.87 | 0.34 | 0.36 ± 0.10 | 81 | 0.76 |
| | spcfy.io | Illumina | $\geq 3$ | 1 | 37.5M | 0.71 | 0.88 | 0.43 | 0.49 ± 0.090 | 68 | 0.62 |
| | mBRAVE | Illumina | $\geq 5$ | 1 | 56.7M | 0.37 | 0.92 | 0.15 | 0.44 ± 0.13 | 85 | 0.77 |
| | MetaWorks | Illumina | $\geq 5$ | 1 | 59.7M | 0.28 | 0.87 | 0.26 | 0.57 ± 0.097 | 73 | 0.56 |
| | QIIME2 | Illumina | $\geq 3$ | 1 | 44.2M | 0.39 | 0.87 | 0.26 | 0.54 ± 0.094 | 72 | 0.58 |
| Stringent filter ($\geq 0.001\%$ of sample reads) | MAP | Nanopore | $\geq 36$ | 1 | 117.8M | 0.65 | 0.90 | 0.47 | 0.30 ± 0.096 | 67 | 0.76 |
| | MAP | Illumina | $\geq 26$ | 1 | 43.7M | 0.66 | 0.90 | 0.33 | 0.35 ± 0.093 | 66 | 0.73 |
| | spcfy.io | Illumina | $\geq 22$ | 1 | 37.5M | 0.66 | 0.89 | 0.42 | 0.45 ± 0.072 | 55 | 0.62 |
| | mBRAVE | Illumina | $\geq 45$ | 1 | 56.6M | 0.77 | 0.90 | 0.35 | 0.34 ± 0.099 | 70 | 0.76 |
| | MetaWorks | Illumina | $\geq 47$ | 1 | 59.5M | 0.78 | 0.87 | 0.34 | 0.49 ± 0.077 | 61 | 0.60 |
| | QIIME2 | Illumina | $\geq 27$ | 1 | 44.1M | 0.79 | 0.87 | 0.34 | 0.48 ± 0.071 | 60 | 0.62 |

502 \*pipelines other than MAP are listed in pre-determined random order

### Validation of MAP with Illumina Data

The approach for validating MAP using Illumina data was identical to that employed for ONT. Biodiversity metrics are in Table 2 while the visualization of Illumina results is presented in Figure S2. While MAP generated accurate biodiversity metrics using Illumina data, they were generally less similar to the barcode data than those derived from ONT. Analyses which downsampled the ONT dataset to the same read count as the Illumina dataset (47.1M) revealed that fewer reads either improved or had little effect on all diversity metrics except for recall (Fig. S3).

### Comparison of MAP to other platforms

Although there was variation among metrics and filter strategies, MAP performed well compared to other platforms (Table 2; Fig. S4–S7). MAP’s strong performance also persisted when only target BINs were used as ground truth (Table S2). In fact, MAP performed consistently well across the three diversity metric categories under all three filter strategies. spcfy.io produced particularly accurate β-diversity estimates while mBRAVE excelled at estimating γ-diversity. These trade-offs reflect differences in reference library usage, filtering philosophy, and OTU clustering approach. MAP’s strong relative performance extended to analyses evaluating the presence/absence of genera or families as reported by the programs themselves (Tables S3 & S4).

MAP’s analysis of the ONT data generated biodiversity metrics with the highest congruence to those obtained from DNA barcode analysis. Considering only Illumina data, while there was variability across metrics and filter strategies, MAP’s performance was comparable to, or better than, the other pipelines. QIIME2 and MetaWorks performed relatively poorly compared to all other pipelines. See Fig. S8 for a visual comparison of β-diversity ordinations, and Fig. S9 for a visual comparison of all metrics with varying proportional filter thresholds.

## DISCUSSION

This paper presents a new bioinformatic pipeline, MAP, developed to facilitate data analysis for metabarcoding studies. While MAP’s support for short-read data is shared by other pipelines like MetaWorks (Porter and Hajibabaei, 2022), mBRAVE (Ratnasingham, 2019), and OptimOTU (Furneaux *et al*., 2025), its ability to analyze PromethION-scale long-read datasets is novel as is its ability to analyze both short– and long-read data. We validated the pipeline through a study of ten bulk arthropod samples and found that MAP returned results that are very similar to DNA barcoding results. MAP also performed well when compared to alternative pipelines. Below, we discuss the key findings of our study and the prospects for MAP.

Because metabarcoding runs can generate many spurious OTUs, stringent filtering is critical for accurate biological inference (Zinger *et al*., 2019; Elbrecht *et al*., 2018). MAP’s denoising approach—combining a background noise threshold derived from negative controls with a replicate consistency test—offers a principled framework for filtering that does not require manual parameter tuning to achieve accurate results. Additional filtering can improve congruence with barcode results, but some valid detections are invariably discarded along with invalid detections when filtering is increased (Zinger *et al*., 2019; Drake *et al*., 2022). MAP was designed to provide the user with flexibility to impose whatever level of filtering they choose, either within-pipeline via the parameters file or post-pipeline by filtering output tables.

MAP’s performance is superior to the other platforms when viewed from the perspective of applicability as it supports the analysis of both long– and short-read data. However, the best choice of pipeline depends on the user’s resources, experience, requirements, and goals. For researchers who lack local compute, spcfy.io and mBRAVE are good options because they are cloud-hosted. Analysis with spcfy.io involves a charge (€25 per sample or less for higher volumes) but it is fast and very user-friendly. mBRAVE is free but it is slower and has a challenging user interface. If local compute is available and open-source analysis is prioritized, MAP is the logical choice for COI barcoding and is essential for long-read data. Among the pipelines that we evaluated, spcfy.io excelled at β-diversity metrics but had relatively poor γ-diversity metrics, a trade-off users should weigh against their project goals. Evaluating the rankings of pipelines across metrics (Table S5) clearly reveals tradeoffs—there is no ‘correct’ answer for any biodiversity metric. However, the present results suggest that the optimal filtration varies among metrics, suggesting that a single dataset should be reanalyzed with different filter strategies to optimize results. Additional high-quality paired barcode-metabarcode datasets must be assembled to establish the generality of these patterns.

The development of an analysis pipeline for full-length COI sequences makes clear the need for a platform to host the results of metabarcoding studies (Bräunig *et al*., 2024). An OTU could be spurious if it is seen for the first time in a particular metabarcoding study, but this probability declines as independent studies detect it. In our study, many high-confidence OTUs gained BINs only after the samples were barcoded, and thus initially represented true dark BINs that were ‘illuminated’ by barcoding (Hausmann *et al*., 2020). Presently, researchers who detect novel OTUs typically submit their data to a sequence repository as raw reads (Buchner *et al*., 2025), and as such they cannot be easily queried.

Making OTUs from metabarcoding studies readily identifiable has clear and immediate benefits. For instance, it is critical to detect species early for efforts to eradicate invasive species (Larson *et al*., 2020). Given the difference in cost, a newly introduced invasive species is far more likely to be detected in a metabarcoded sample than as a vouchered barcoded specimen. Moreover, a sample uploaded to a database could be flagged as a high-value sample for barcode analysis if it included many novel OTUs, suggesting it would yield a high number of new BINs. The development of a high-quality centralized database for metabarcoding OTUs should be a high priority for the metabarcoding community.

In conclusion, MAP enables researchers to generate high-quality, readily interpretable results from multiplexed metabarcoding sequencing runs. It additionally facilitates the production and eventual preservation of high-quality OTUs from metabarcoding studies, with outputs addressing both basic and applied questions about biodiversity. While MAP currently operates from the command line and targets users with bioinformatics expertise, it will soon be implemented as the back end of ONTOLOGY™, a user-friendly software platform that will make metabarcoding analysis accessible to non-specialists.

### Data accessibility

Demultiplexed Oxford Nanopore Technologies and Illumina sequences are available at the NCBI SRA BioProject Accession PRJNA1496812. Scripts and other files used for MAP and benchmarking analysis provided at Figshare (doi: 10.6084/m9.figshare.33027503). MAP package can be accessed and pulled from https://github.com/cbg-innov/MAP/pkgs/container/map, with additional information for the package available from the MAP GitHub repository: https://github.com/cbg-innov/MAP.

### Author contributions

- Fieldwork and Sample Collection: PDNH
- Data collection and curation: RMF, SJ, SWJP
- Writing first draft: KAT & PDNH
- Revision: All authors
- Software Implementation: SWJP, NB
- Software Design: SWJP, NB, KAT, SP, EO, PDNH
- Data Analysis: KAT

### Ethics statement

This manuscript complies with all relevant policies concerning ethics in research.

## Acknowledgements

We thank Jayme Sones, Jaclyn McKeown, Madison Gardiner, Hikaru Morashita, Victoria Smelko in the CBG’s Specimens Unit for databasing, imaging, and curating the voucher specimens. We thank Suresh Naik for performing hit-picking while Amy Thompson oversaw DNA extraction, which was performed by the CBG’s Genomics Unit. The CBG Genomics unit also performed some of the PCRs.

## Use of AI

AI tools (including Anthropic’s Claude Code) were used to assist with software development, data analysis, and manuscript editing.

## SUPPLEMENTARY MATERIALS

### SUPPLEMENTARY METHODS

#### COI-5P Error correction

##### Automated homopolymer correction script

As nanopore sequencing has a high error rate in homopolymer runs, MAP incorporates an automated correction script (homopolymer_error_fix.py) which screens for indel errors in homopolymer tracts. Each OTU sequence from a run is searched using vsearch against a known, validated reference set of sequences trimmed to the exact barcode region (details for the construction of this set are given below) to find its nearest neighbour. The sequence and its neighbour are aligned, and end-correction— where the query sequence is either trimmed or padded with Ns to the exact length of neighbour sequence—is carried out.

Next, the sequence is searched for any internal gaps. If present, the following logic is applied:

- Only one or two consecutive gaps or insertions are corrected;
- Only corrected if they are part of, or adjacent to, a run of the same nucleotide (by default, a minimum of 4);
- If gaps are in the query sequence, they are replaced with a lowercase ‘n’ (to distinguish from uppercase ‘N’ which may result from ambiguities in consensus calling so users can easily recognize when automatic correction has been applied);
- If gaps are in the reference sequence, a corresponding number of letters from the homopolymer are deleted from the query sequence.

Sequences are output to a fasta file (“edited_seqs.fasta”). If automatic correction was carried out, the text “|autoedit” is appended to the fasta header sequence. The correction script also screens for stop codons and HMM issues which may remain, using the codon translation table derived from the nearest neighbour (these indicate either errors which could not be corrected using the homopolymer logic, or likely NUMTs). In these cases, “|STOP” and/or “|indel” are also added to the fasta header. Finally, if residual one– or two-base indels remain in the alignment which are not associated with homopolymers, the script will attempt to edit these by replacing gaps with “n” or deleting extra bases, and test whether these edits eliminate HMM issues. If so, these edits are not output automatically, but the text “|check” is appended to the fasta header in the output, and users may investigate whether they consider these to be sequencing errors by examining the associated raw reads. For each OTU sequence where any edits were made or suggested, an individual alignment is output to a folder (“alignment_dir”) containing the nearest neighbour match used as the reference, the original unedited sequence, and any edited version(s).

##### Reference Panel creation

The error correction script relies upon a reference dataset of true COI sequences correctly trimmed to the exact barcode region and containing no indel errors. It was generated by subsampling the BOLDistilled reference library which compresses all available taxonomic diversity of COI sequences in BOLD into a minimal number of sequences. A table was created from all sequences in the current BOLDistilled set with the following columns: ProcessID, nucleotide sequence, translated amino acid sequence, and translation table identifier (e.g., “5” for invertebrate mitochondrial). This table was filtered using a custom Python script (hp_error_ref_set_process.py) as follows:

- All rows where the nucleotide sequence contained an “N” character (ambiguous nucleotide) were dropped.
- All rows where the amino acid sequence contained either “*” (stop codon) or “X” (unknown amino acid) were dropped.
- Duplicate amino acid sequences were removed; i.e., only one row representative of each unique amino acid was retained.
- The remaining amino acid sequences were then parsed using a regular expression which matches the 5’ and 3’ ends of the barcode region. Only those rows matching the specified pattern were kept.
- The nucleotide sequences of the remaining rows were then trimmed to the exact region corresponding to the amino acid pattern above.
- Finally, all nucleotide sequences with an overall length outside 640-670bp, or whose length is not divisible by 3, were deleted.
- The remaining nucleotide sequences were output as a fasta file in which each sequence header contains the ProcessID and mitochondrial translation table of the source organism.

Following this procedure, ∼1.5 million sequences in the full BOLDistilled set were reduced to ∼330,000, allowing most query sequences to rapidly find a valid nearest-neighbour sequence to align against, thus enabling the error correction script to determine the correct reading frame and to identify any indels which are potential sequencing errors. The correct codon translation table is also extracted from the fasta header of the nearest neighbour and used to check for stop codons in the query sequence.

#### Metabarcoding

DNA extraction for metabarcoding analysis involved overnight lysis at 56 °C in a buffer consisting of 700 mM guanidine thiocyanate (MilliporeSigma), 30 mM Tris-HCl pH 8.0 (MilliporeSigma), 30 mM EDTA pH 8.0 (Thermo Fisher Scientific), 5% Tween-20 (MilliporeSigma), and 0.5% Triton X-100 (MilliporeSigma). The resultant lysate was separated from the specimens by filtration through a Microfunnel 0.2 μm Supor Membrane Filter (Pall Life Sciences) into a sterile container. The specimens were placed in fresh 95% ethanol for subsequent barcode analysis (see ‘DNA BARCODING’ below). DNA was extracted from eight aliquots of the lysate from each sample, creating a total of 80 DNA extracts. DNA extracts were generated using a standard glass fiber protocol (Ivanova *et al*., 2006). PCR was performed in a 96-well plate containing all 80 DNA extracts plus eight positive and eight negative control wells. Each 12.5 μL PCR reaction consisted of 2 μL of DNA plus 10 μL of master mix: 5% trehalose (Fluka Analytical), 1× PlatinumTaq buffer (Thermo Fisher Scientific), 2.5 mM MgCl_2_ (Thermo Scientific), 0.05 mM dNTPs (KAPA Biosystems), 0.1 μM of each indexed primer (IDT; Table S1), 0.3 U PlatinumTaq (Thermo Fisher Scientific), and molecular grade water (Hyclone, Thermo Fisher). The thermocycling regime was 94 °C for 2 min, five cycles of [94 °C for 40 s, 45 °C for 40 s, 72 °C for 1 min], 25 cycles of [94 °C for 40 s, 51 °C for 40 s, 72 °C for 1 min], and a final extension of 72 °C for 2 min.

Amplicons were prepared for sequencing using Oxford Nanopore Technologies’ (ONT) Ligation Sequencing Kit (SQL-LSK114) following the manufacturer’s protocol. The final library was sequenced using a PromethION Flow Cell (FLO-PRO114M) on a P2 Solo. During sequencing, live base-calling was enabled (Dorado v7.4.14) using the SUP—super accurate—base-calling model. The run generated 179,806,945 base-called reads.

These reads were analyzed with MAP v1.0.1 using the parameters file in the online data repository. Analysis on a computer provisioned with an AMD Ryzen ThreadRipper 7980X 128-core CPU, 128 GB RAM, an NVIDIA GeForce RTX 4090 GPU, and running Ubuntu 24.04.1 LTS required 36 hr. All OTUs that survived denoising with at least two reads were retained. This threshold is lower than most users might select, but we use it here for illustrative purposes.

#### DNA Barcoding

The specimens in each bulk sample were placed into a 10 cm Petri dish for processing. Each small specimen (< 3 mm) was placed as a whole voucher into one of the wells in a 96-well plate containing 5 ul of 95% ethanol and then photographed. Larger specimens (> 3 mm) were pinned and then photographed. DNA was extracted from whole small specimens, or from a leg segment from pinned specimens using a magnetic bead-based method (solid-phase reversible immobilization [SPRI]; Ivanova et al. [in prep]).

The 658 bp COI barcode region was amplified using the same primer sequences used for metabarcoding (for UMI details, refer to the ‘10K’ data set in Hebert et al. (2024)). Reaction conditions were identical to those used for metabarcoding, and thermocycling was identical except that 45 cycles—rather than 25 cycles—were used for the second phase of amplification. All PCR products were pooled, and library preparation was carried out using the ONT Ligation Sequencing Kit (SQK-LSK114) following a published protocol (Floyd *et al*., 2023). Libraries were then sequenced on either an ONT MinION Mk-1b or Mk-1d sequencer.

Because the results from DNA barcoding provide the definitive reference data on the identity and number of specimens of each BIN in each bulk sample, it was important to maximize barcode recovery. To that end, we conducted four sequencing runs for barcoding. The first sequencing run (n = 7,044 specimens), which was conducted on a MinION flow cell, omitted many of the springtails and ants that were likely to belong to few species. A second run (n = 4,465 specimens), which also used a MinION flow cell, analyzed all the latter specimens plus some other very common morphospecies from the samples. A third run, which employed a Flongle Flow Cell, reanalyzed all DNA extracts that did not return a DNA sequence from the first two sequencing runs (n = 870 specimens). The final run repeated PCR followed by sequencing on a Flongle for all specimens that did not generate a DNA sequence after two prior attempts (n = 362 specimens). Specimens that generated a sequence that was not barcode compliant— for example those with a stop codon—were not reanalyzed. Each of these DNA extracts was PCR-amplified twice—once after a 50× dilution of the DNA extract, and another time with 60 PCR cycles. DNA extraction was not re-attempted for any specimens.

All barcode results were uploaded to BOLD and are accessible as a public dataset DS-PHAUS24 (dx.doi.org/10.5883/DS-PHAUS24). Note that the BOLD data portal filters out low-quality records, leading to lower numbers of specimens than in the dataset when accessed via the BOLD workbench. Since MAP validation required direct comparison of BIN assignments between the barcoding and metabarcoding results, we opted to assign BINs and taxonomy to the barcode results via MAP’s BIN matching module using the April 2026 COI BOLDistilled library. In this way, taxonomy was assigned to sequences using identical methods for both datasets. Note also that this leads to minor differences in BIN counts and assignments in this manuscript compared to what is in the BOLD dataset.

#### Benchmarking

Benchmarking could only employ the Illumina paired-end dataset. Attempts to process a PromethION-scale dataset using PRONAME (Dubois *et al*., 2024) failed due to the high space and memory requirements.

##### MAP

Default parameters were used for ONT dataset (3 minimum reads per OTU). With the Illumina dataset, the following deviations were applied, in order to compensate for the lower error rate associated with Illumina paired end sequencing relative to ONT (i.e., the defaults supplied in MAP.sh script/README and in provided parameters.xlsx file): high_dens_prop =0.15, UMI_overlap_min=1.0, and 2 minimum reads per OTU. The remaining deviations were to accommodate for the shorter UMI sequences and comprise the parameter settings: error_umi1 (and 2) = 0, umi_overlap = 1, and a maximum intra-sample OTU clustering threshold of 2.5%.

##### spcfy.io

Metabarcode data was uploaded to spcfy.io and analyzed using all default settings and employing the Animal COI reference libraries and SPARK output.

##### mBRAVE

We compared the output of MAP to mBRAVE using the parameters for COI listed in Table S6. We used the following reference libraries in the following order: SYS-MBRAVEC, SYS-CRLDIPTERA, SYS-CRLNONDIPINSECTA, SYS-CRLNONINSECTARTH, SYS-CRLNONARTHINVERT.

##### MetaWorks

We first ran the MetaWorks ESV pipeline using default parameters except for a minimum read count of 2. The barcode was set as ‘COI’. We allowed for pseudogene filtering using the provided BOLD hidden Markov model (HMM), targeting ‘Arthropoda’. Afterward, we ran the OTU pipeline, using the RDP COI rRNAClassifier.properties file v5.1.0. Configuration files are available in the code repository associated with this manuscript (See <u>Data & Code Accessibility)</u>.

##### QIIME2

The QIIME2 metabarcoding workflow was run on demultiplexed .fastq files that were initially merged, primer trimmed, and quality filtered by MetaWorks. Denoising was run using a memory-modified version of Dada2 denoise-single (see, after dropping contigs shorter than 388 bp and longer than 430 bp, respectively. The denoised ASVs were clustered to OTUs at 97% identity and classified using global alignment with a 60% identity threshold. Command line prompts are available in the code repository associated with this article (See <u>Data & Code Accessibility).</u>

## SUPPLEMENTARY FIGURES

**Fig. S1.**
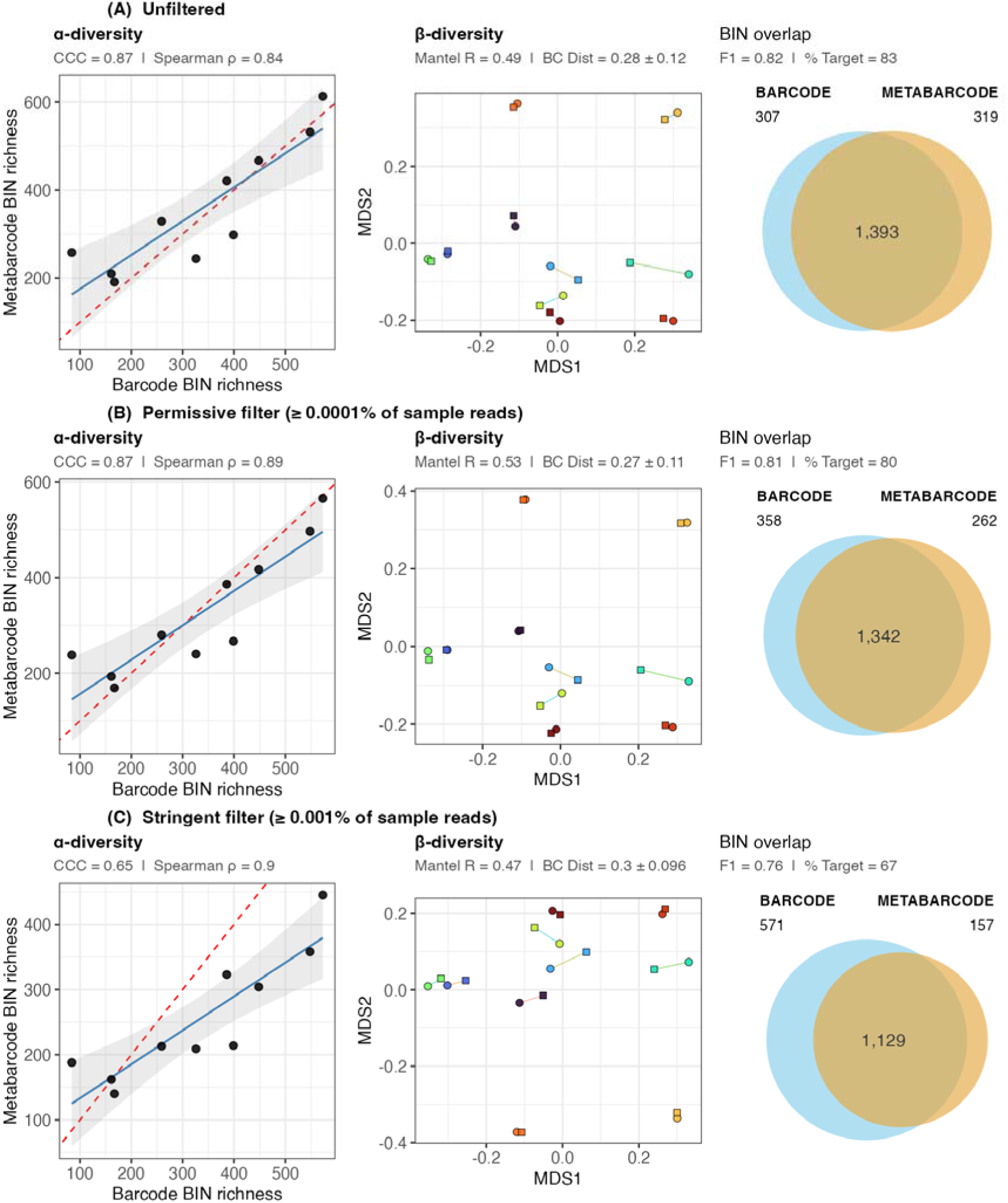
Impact of filtering regimes on biodiversity metrics based on MAP analysis of ONT data from 10 bulk samples of arthropods. Data presented in Fig. 2 of the main text are presented again to facilitate comparison. In β-diversity plots, circles are barcode and squares are metabarcode. Perfect matches between the datasets yield a value of ‘1’ for all metrics except for BC distance, where a perfect match yields a value of ‘0’. See Fig. 2 caption of main text for more details.

**Fig. S2.**
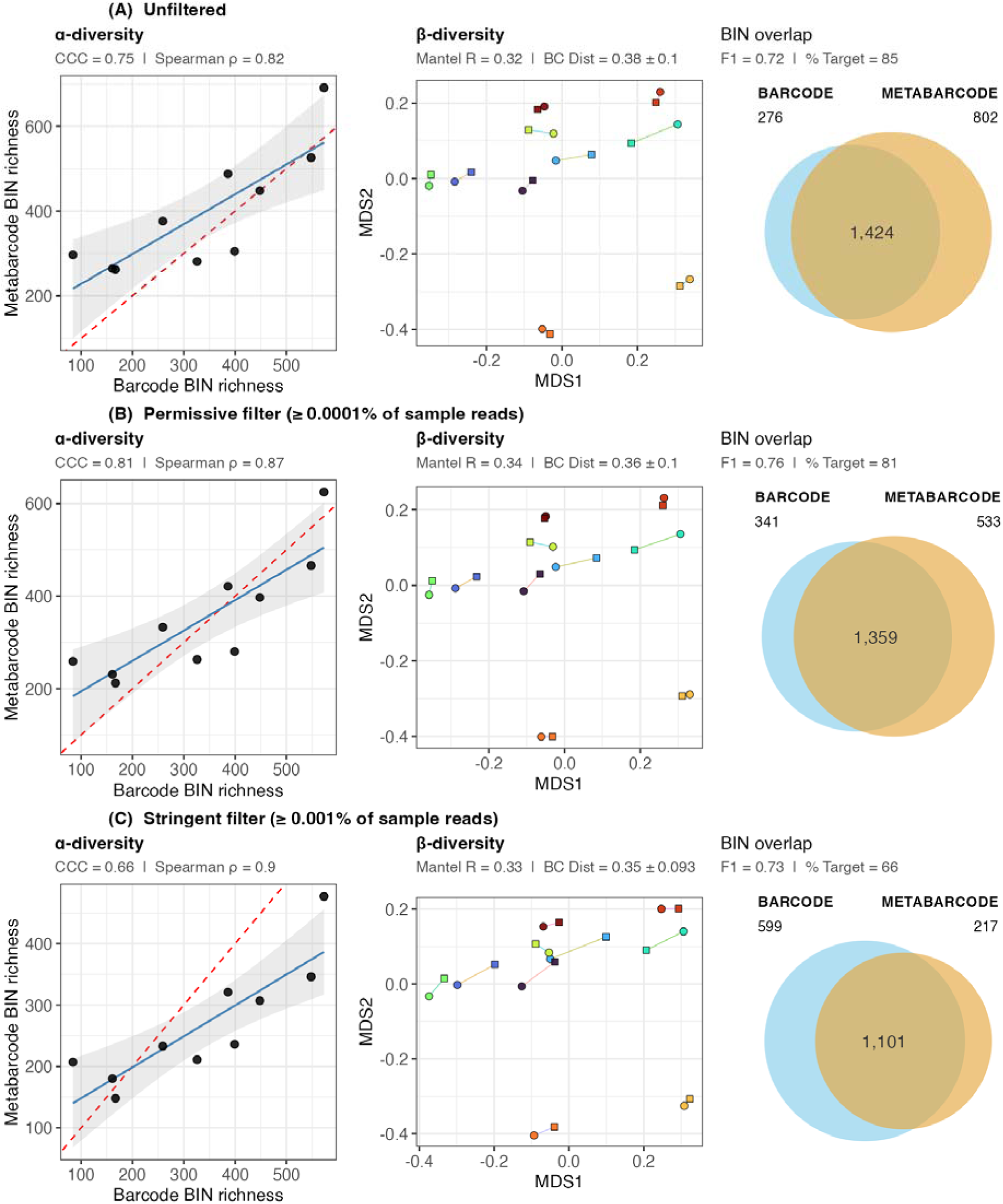
Impact of filtering regimes on biodiversity metrics based on MAP analysis of Illumina data from 10 bulk samples of arthropods. In β-diversity plots, circles are barcode and squares are metabarcode. Perfect matches between the datasets yield a value of ‘1’ for all metrics except for BC distance, where a perfect match yields a value of ‘0’. See Fig. 2 caption of main text for more details.

**Fig. S3.**
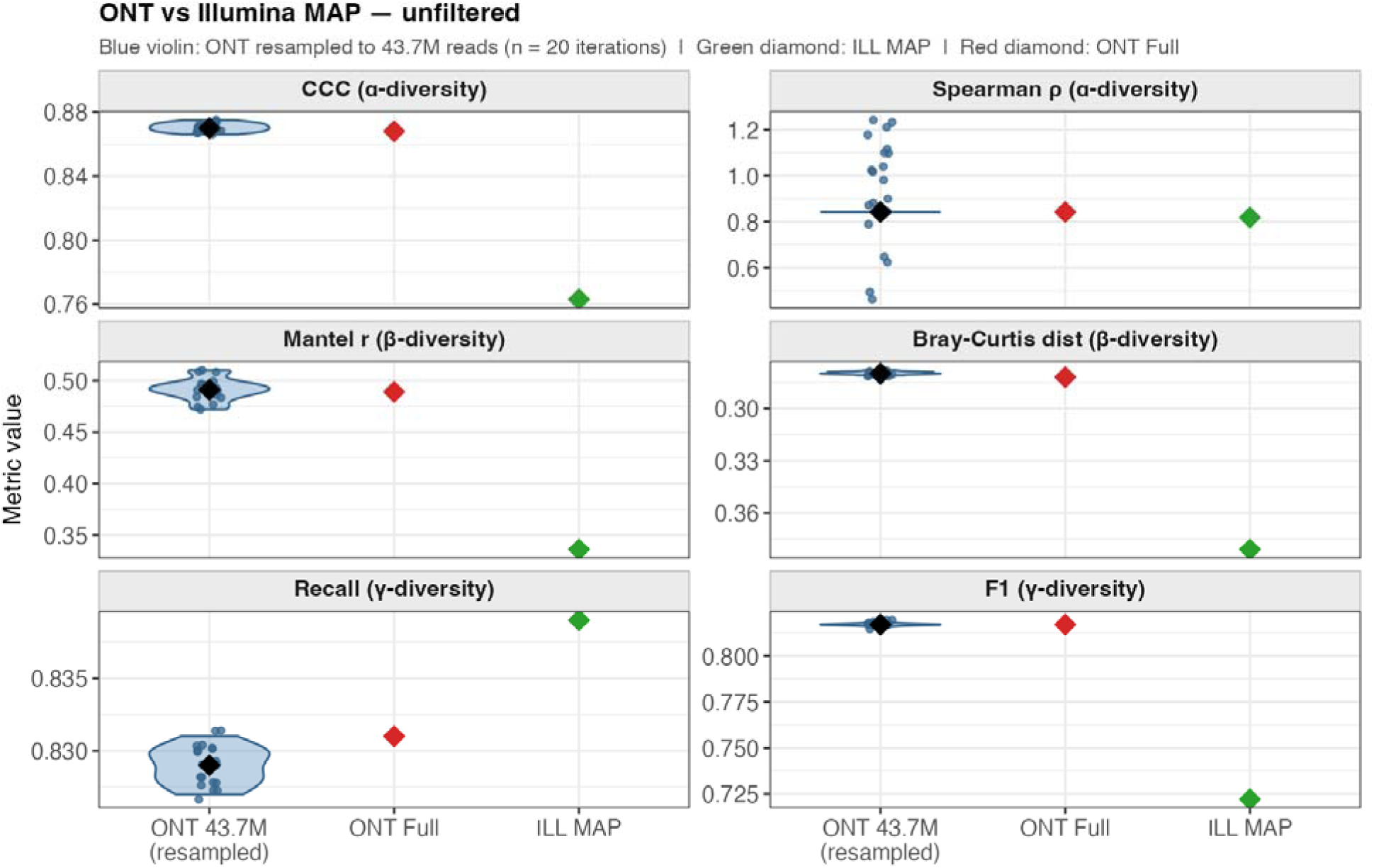
Analysis of downsampled ONT data. The ONT data was resampled to 47.1M reads 20 times and diversity metrics were re-calculated; median is shown in black. Higher values are better for all metrics except Bray-Curtis distance. The y-axis is inverted in the plot of Bray-Curtis distance to facilitate visual comparison.

**Fig. S4.**
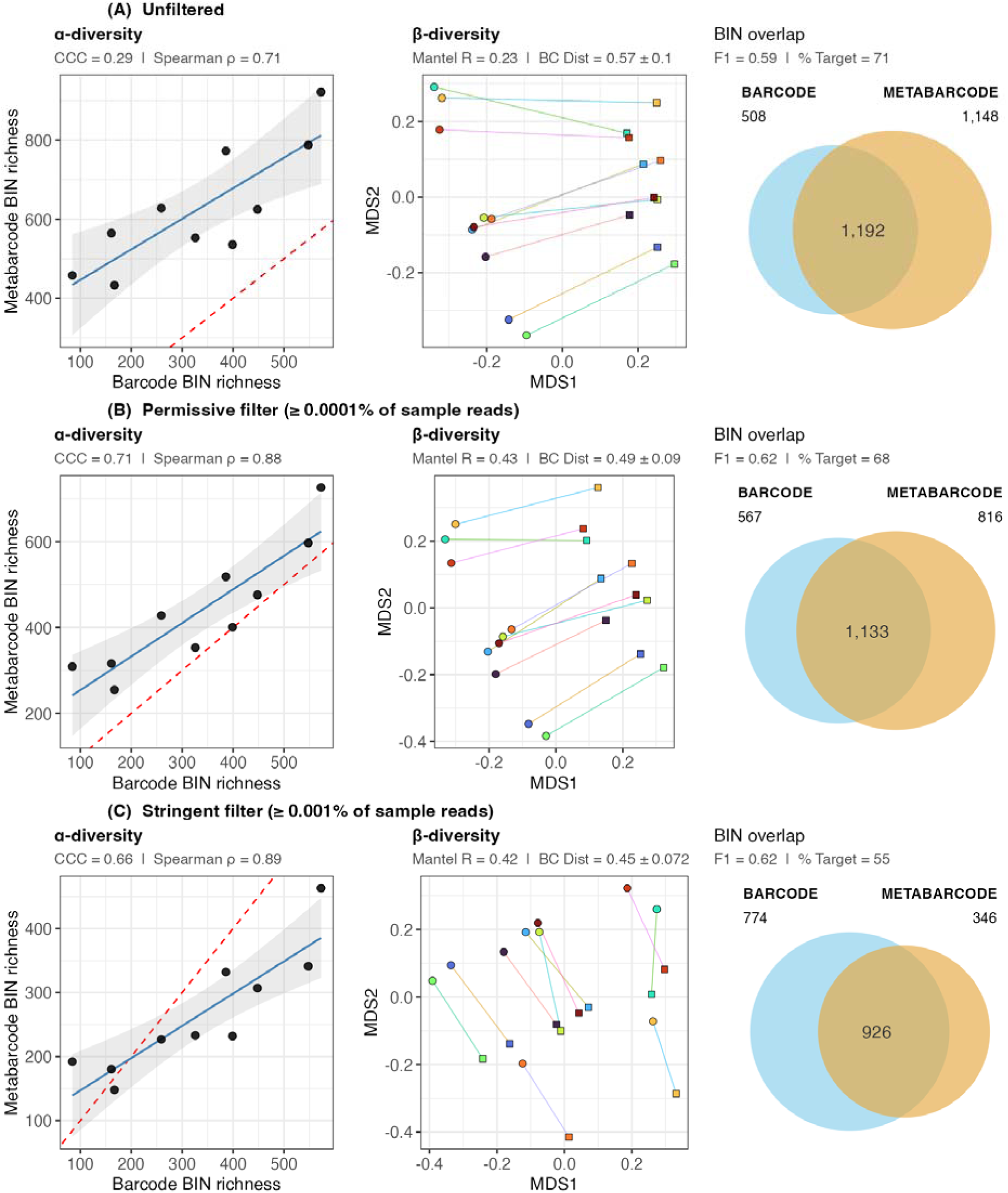
Impact of filtering regimes on biodiversity metrics based on spcfy.io analysis of Illumina data from 10 bulk samples of arthropods. In β-diversity plots, circles are barcode and squares are metabarcode. Perfect matches between the datasets yield a value of ‘1’ for all metrics except for BC distance, where a perfect match yields a value of ‘0’. See Fig. 2 caption of main text for more details.

**Fig. S5.**
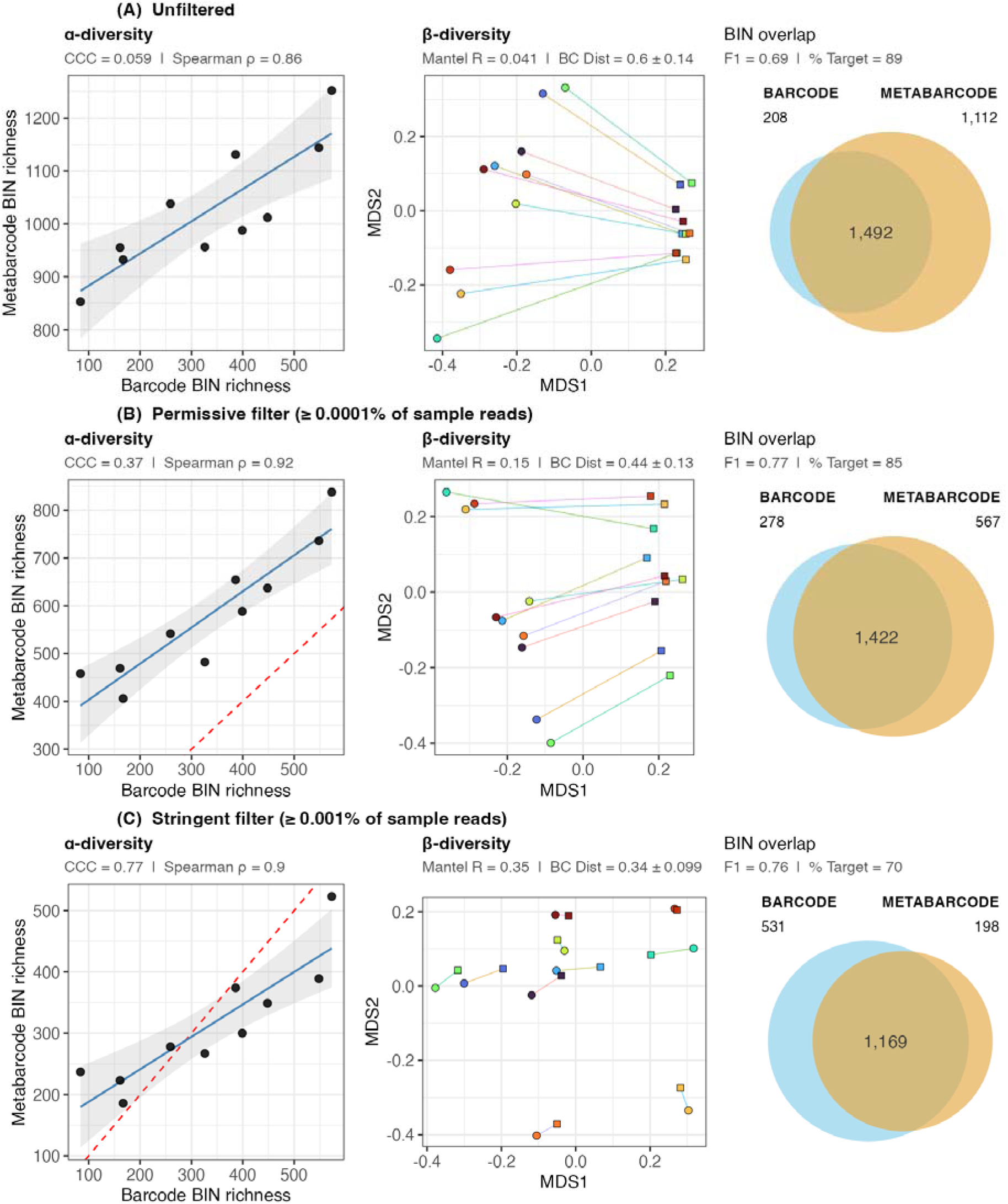
Impact of filtering regimes on biodiversity metrics based on mBRAVE analysis of Illumina data from 10 bulk samples of arthropods. In β-diversity plots, circles are barcode and squares are metabarcode. Perfect matches between the datasets yield a value of ‘1’ for all metrics except for BC distance, where a perfect match yields a value of ‘0’. See Fig. 2 caption of main text for more details. Note that the dashed line is not visible in (A) due to the points being far above the 1:1 line.

**Fig. S6.**
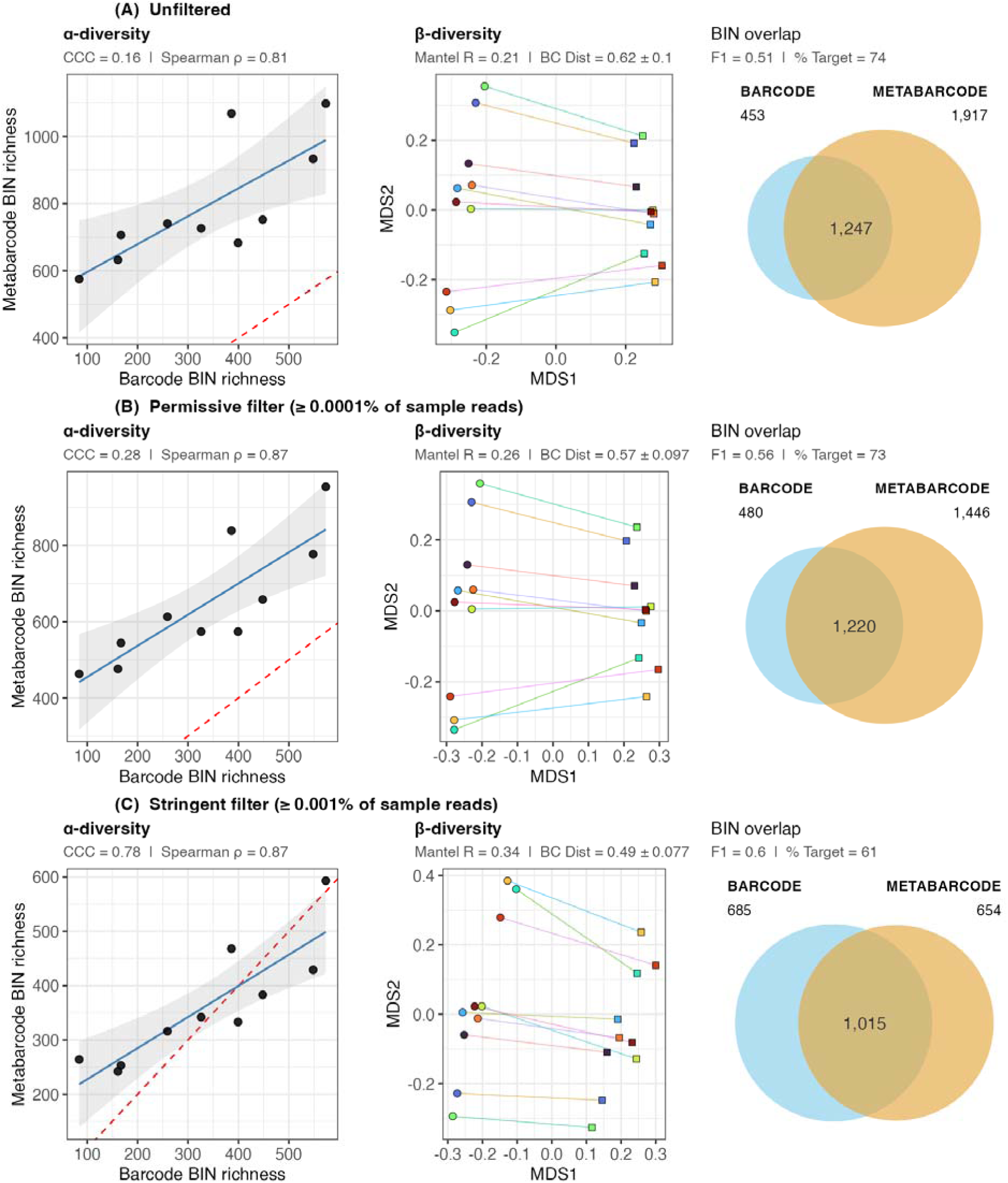
Impact of filtering regimes on biodiversity metrics based on MetaWorks analysis of Illumina data from 10 bulk samples of arthropods. In β-diversity plots, circles are barcode and squares are metabarcode. Perfect matches between the datasets yield a value of ‘1’ for all metrics except for BC distance, where a perfect match yields a value of ‘0’. See Fig. 2 caption of main text for more details.

**Fig. S7.**
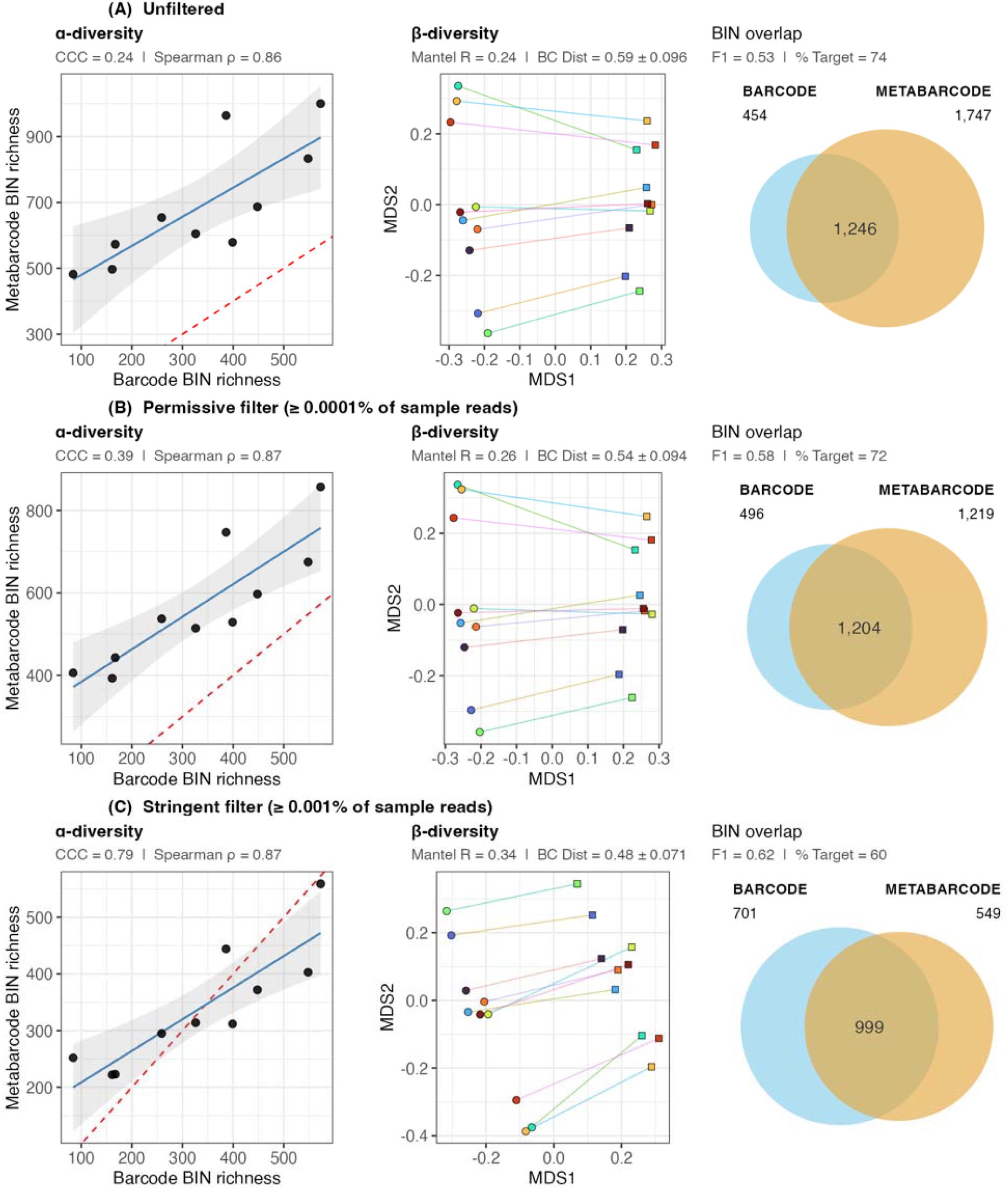
Impact of filtering regimes on biodiversity metrics based on QIIME2 analysis of Illumina data from 10 bulk samples of arthropods. In β-diversity plots, circles are barcode and squares are metabarcode. Perfect matches between the datasets yield a value of ‘1’ for all metrics except for BC distance, where a perfect match yields a value of ‘0’. See Fig. 2 caption of main text for more details.

**Fig. S8.**
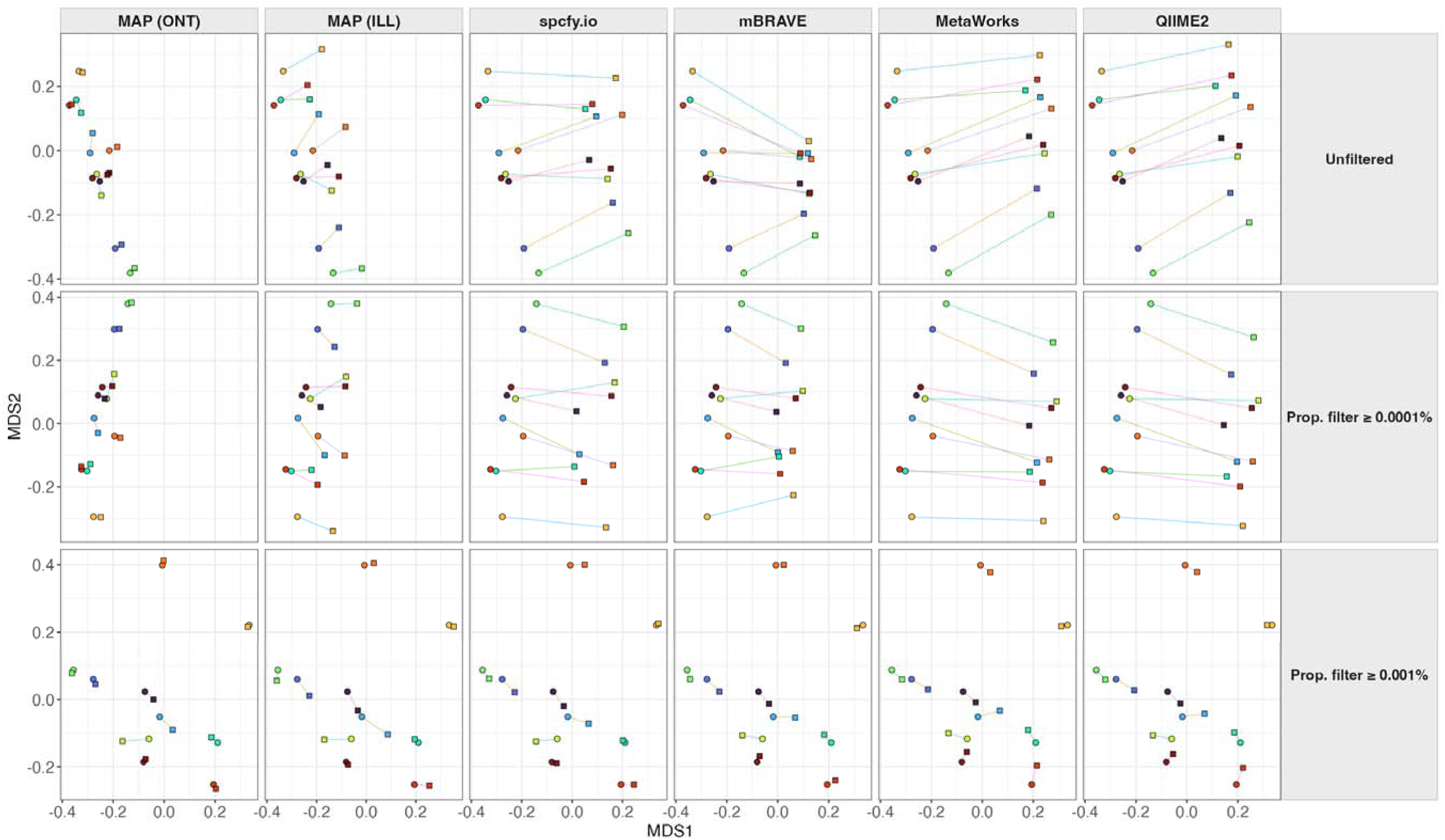
β-diversity ordination plots for the three filtration strategies. The axes are identical within rows.

**Fig. S9.**
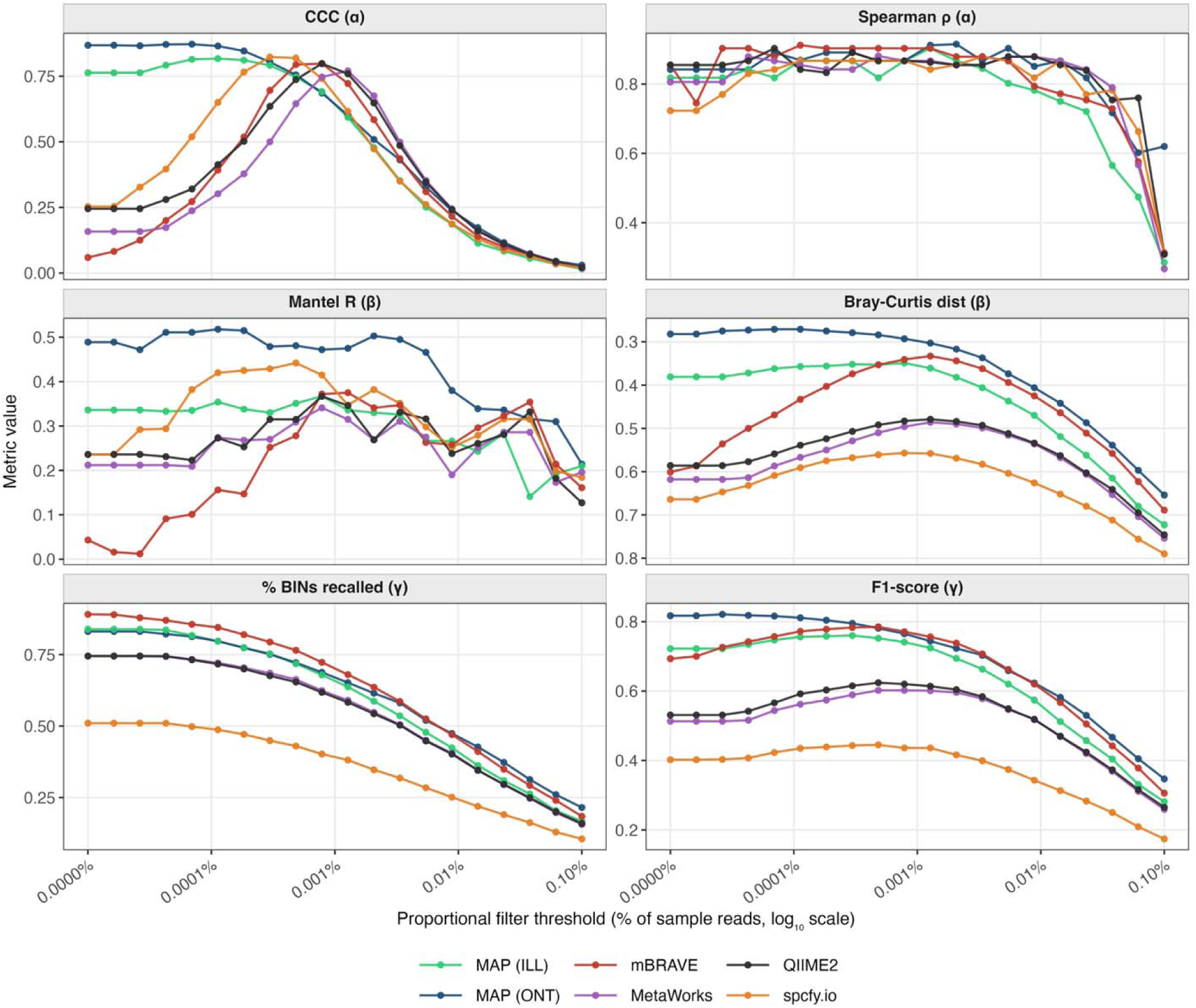
All six diversity metrics at various filter thresholds. 0% is unfiltered. Note that the y-axis is inverted in the Bray-Curtis distance panel to facilitate visual comparison. Values used in Table 2 and similar tables are 0.0001% and 0.001%.

## SUPPLEMENTARY TABLES

**Table S1:**
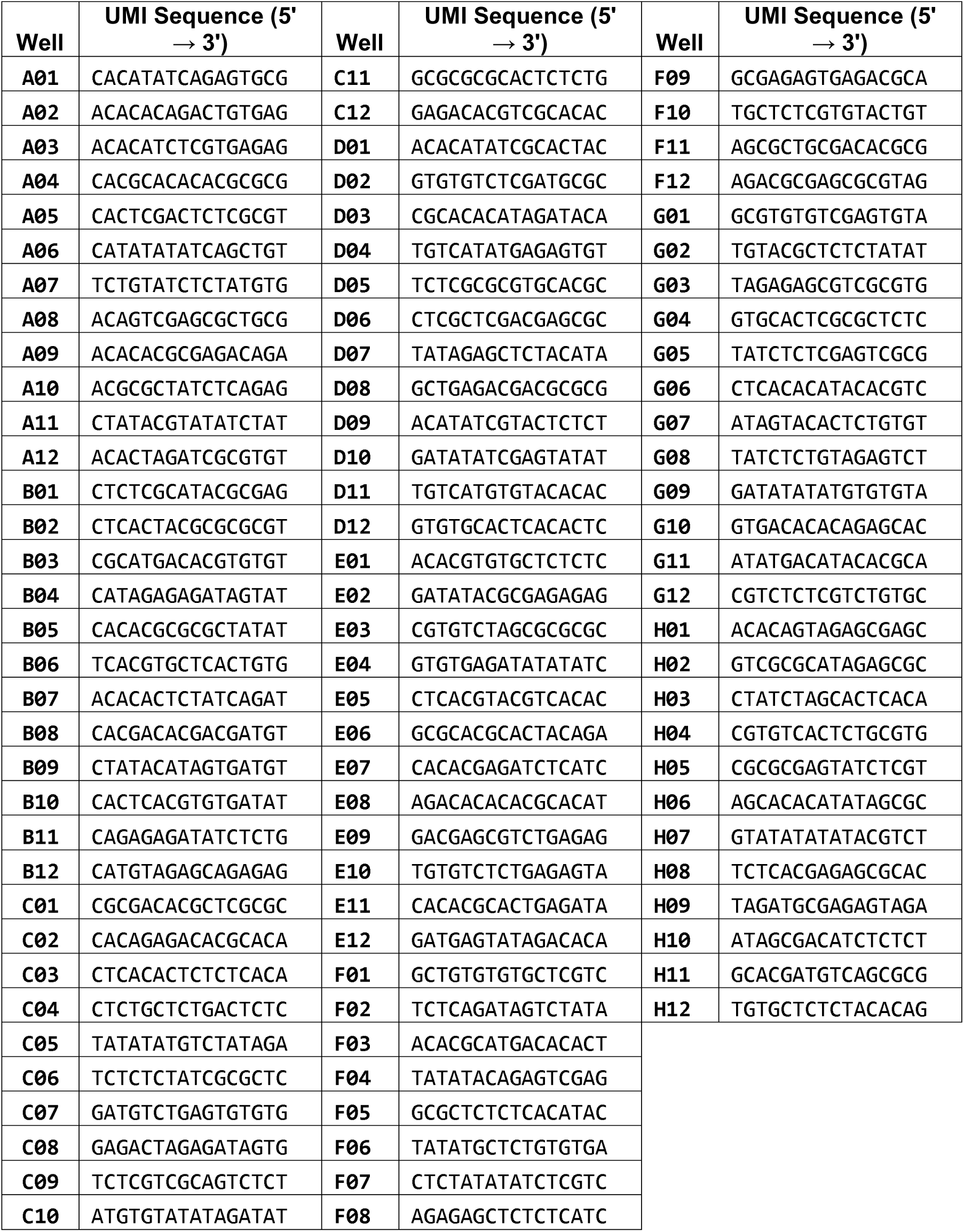
Symmetrical UMI sequences by plate well. All PCRs contained the same COI primers (C_LepFolF/C_LepFolR) tailed with unique well-specific UMIs. Forward and reverse primers were symmetrically tailed with the same UMI sequence.

**Table S2.** Summary biodiversity metrics computed for target barcodes only, rather than Target +NTS; darker green indicates better performance within each metric and filter strategy.

| Filter Strategy | Software | Dataset | Min. Reads | Min. Reps | # Reads in OTUs | CCC ($\alpha$) | Spearman $\rho$ ($\alpha$) | Mantel $R$ ($\beta$) | BC Dist ± SD ($\beta$) | % Target ($\gamma$) | F1-score ($\gamma$) |
| --- | --- | --- | --- | --- | --- | --- | --- | --- | --- | --- | --- |
| Unfiltered | MAP | Nanopore | 3 | 1 | 118.1M | 0.84 | 0.84 | 0.39 | 0.29 ± 0.15 | 83 | 0.81 |
| | MAP | Illumina | 2 | 1 | 43.8M | 0.71 | 0.82 | 0.25 | 0.39 ± 0.13 | 85 | 0.72 |
| | spcfy.io | Illumina | 1 | 1 | 37.5M | 0.26 | 0.71 | 0.23 | 0.59 ± 0.11 | 71 | 0.59 |
| | mBRAVE | Illumina | 1 | 1 | 56.7M | 0.056 | 0.85 | 0.022 | 0.62 ± 0.14 | 89 | 0.69 |
| | MetaWorks | Illumina | 3 | 1 | 61.9M | 0.13 | 0.72 | 0.11 | 0.64 ± 0.11 | 74 | 0.49 |
| | QIIME2 | Illumina | 2 | 1 | 44.2M | 0.23 | 0.85 | 0.19 | 0.60 ± 0.11 | 74 | 0.53 |
| Permissive filter ($\geq 0.0001\%$ of sample reads) | MAP | Nanopore | $\geq 4$ | 1 | 117.9M | 0.87 | 0.89 | 0.43 | 0.28 ± 0.15 | 80 | 0.81 |
| | MAP | Illumina | $\geq 3$ | 1 | 43.8M | 0.80 | 0.87 | 0.27 | 0.37 ± 0.13 | 81 | 0.75 |
| | spcfy.io | Illumina | $\geq 3$ | 1 | 37.5M | 0.66 | 0.88 | 0.41 | 0.50 ± 0.11 | 68 | 0.62 |
| | mBRAVE | Illumina | $\geq 5$ | 1 | 56.7M | 0.33 | 0.92 | 0.10 | 0.46 ± 0.15 | 85 | 0.77 |
| | MetaWorks | Illumina | $\geq 5$ | 1 | 61.9M | 0.23 | 0.87 | 0.18 | 0.60 ± 0.11 | 73 | 0.54 |
| | QIIME2 | Illumina | $\geq 3$ | 1 | 44.2M | 0.35 | 0.87 | 0.21 | 0.56 ± 0.11 | 72 | 0.58 |
| Stringent filter ($\geq 0.001\%$ of sample reads) | MAP | Nanopore | $\geq 36$ | 1 | 117.8M | 0.71 | 0.90 | 0.39 | 0.30 ± 0.13 | 67 | 0.75 |
| | MAP | Illumina | $\geq 26$ | 1 | 43.7M | 0.71 | 0.90 | 0.26 | 0.36 ± 0.13 | 66 | 0.73 |
| | spcfy.io | Illumina | $\geq 22$ | 1 | 37.5M | 0.72 | 0.89 | 0.34 | 0.45 ± 0.099 | 55 | 0.62 |
| | mBRAVE | Illumina | $\geq 45$ | 1 | 56.6M | 0.79 | 0.90 | 0.27 | 0.35 ± 0.13 | 70 | 0.76 |
| | MetaWorks | Illumina | $\geq 47$ | 1 | 61.6M | 0.75 | 0.87 | 0.26 | 0.50 ± 0.099 | 60 | 0.59 |
| | QIIME2 | Illumina | $\geq 27$ | 1 | 44.1M | 0.79 | 0.87 | 0.28 | 0.49 ± 0.097 | 60 | 0.61 |

**Table S3.**
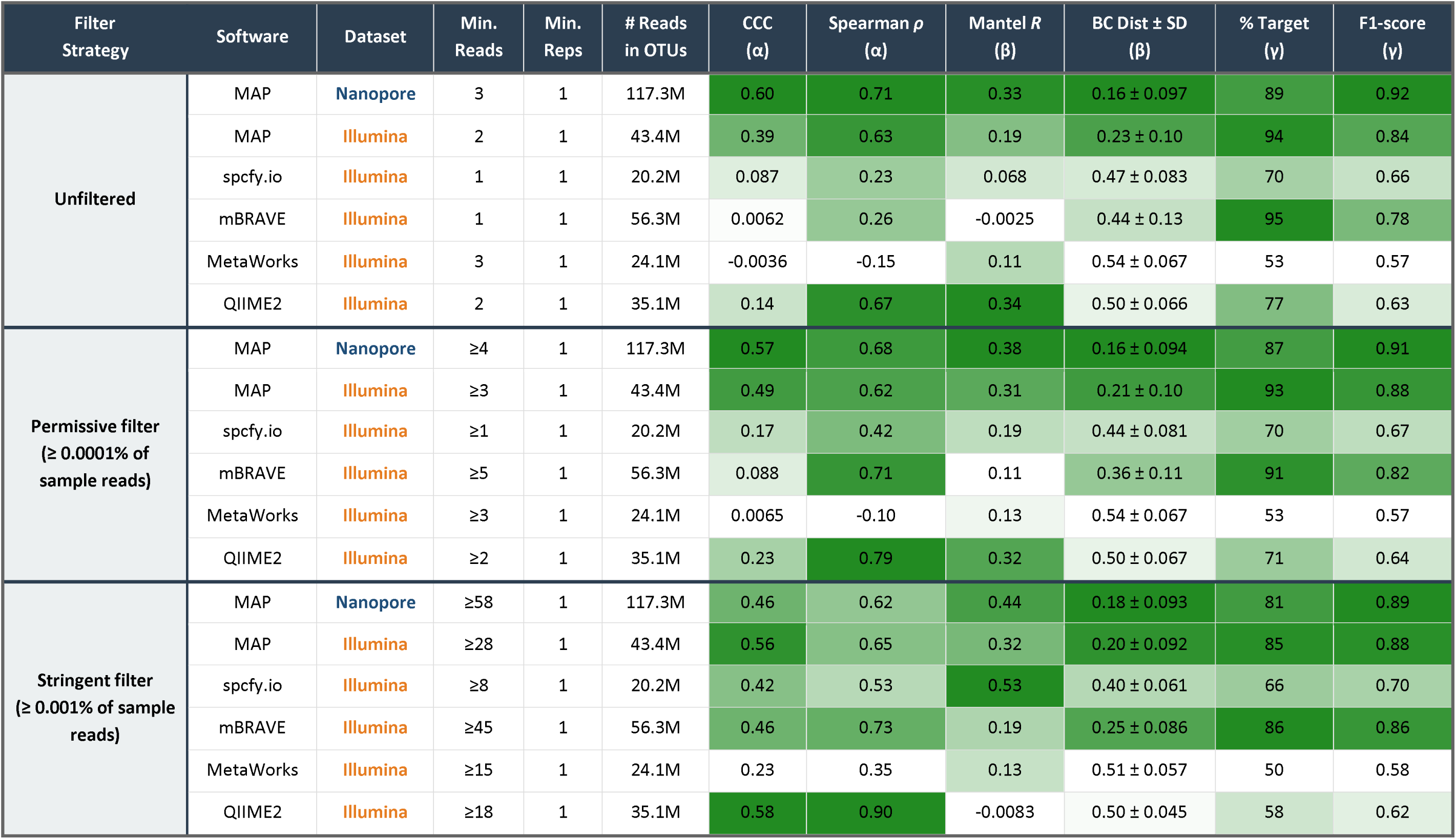
Summary biodiversity metrics across filter strategies using the Target + NTS dataset with <u>Family</u> as the unit of analysis; darker green indicates better performance within each metric and filter strategy.

| Filter Strategy | Software | Dataset | Min. Reads | Min. Reps | # Reads in OTUs | CCC ($\alpha$) | Spearman $\rho$ ($\alpha$) | Mantel $R$ ($\beta$) | BC Dist ± SD ($\beta$) | % Target ($\gamma$) | F1-score ($\gamma$) |
| --- | --- | --- | --- | --- | --- | --- | --- | --- | --- | --- | --- |
| Unfiltered | MAP | Nanopore | 3 | 1 | 117.3M | 0.60 | 0.71 | 0.33 | 0.16 ± 0.097 | 89 | 0.92 |
| | MAP | Illumina | 2 | 1 | 43.4M | 0.39 | 0.63 | 0.19 | 0.23 ± 0.10 | 94 | 0.84 |
| | spcfy.io | Illumina | 1 | 1 | 20.2M | 0.087 | 0.23 | 0.068 | 0.47 ± 0.083 | 70 | 0.66 |
| | mBRAVE | Illumina | 1 | 1 | 56.3M | 0.0062 | 0.26 | -0.0025 | 0.44 ± 0.13 | 95 | 0.78 |
| | MetaWorks | Illumina | 3 | 1 | 24.1M | -0.0036 | -0.15 | 0.11 | 0.54 ± 0.067 | 53 | 0.57 |
| | QIIME2 | Illumina | 2 | 1 | 35.1M | 0.14 | 0.67 | 0.34 | 0.50 ± 0.066 | 77 | 0.63 |
| Permissive filter ($\geq 0.0001\%$ of sample reads) | MAP | Nanopore | $\geq 4$ | 1 | 117.3M | 0.57 | 0.68 | 0.38 | 0.16 ± 0.094 | 87 | 0.91 |
| | MAP | Illumina | $\geq 3$ | 1 | 43.4M | 0.49 | 0.62 | 0.31 | 0.21 ± 0.10 | 93 | 0.88 |
| | spcfy.io | Illumina | $\geq 1$ | 1 | 20.2M | 0.17 | 0.42 | 0.19 | 0.44 ± 0.081 | 70 | 0.67 |
| | mBRAVE | Illumina | $\geq 5$ | 1 | 56.3M | 0.088 | 0.71 | 0.11 | 0.36 ± 0.11 | 91 | 0.82 |
| | MetaWorks | Illumina | $\geq 3$ | 1 | 24.1M | 0.0065 | -0.10 | 0.13 | 0.54 ± 0.067 | 53 | 0.57 |
| | QIIME2 | Illumina | $\geq 2$ | 1 | 35.1M | 0.23 | 0.79 | 0.32 | 0.50 ± 0.067 | 71 | 0.64 |
| Stringent filter ($\geq 0.001\%$ of sample reads) | MAP | Nanopore | $\geq 58$ | 1 | 117.3M | 0.46 | 0.62 | 0.44 | 0.18 ± 0.093 | 81 | 0.89 |
| | MAP | Illumina | $\geq 28$ | 1 | 43.4M | 0.56 | 0.65 | 0.32 | 0.20 ± 0.092 | 85 | 0.88 |
| | spcfy.io | Illumina | $\geq 8$ | 1 | 20.2M | 0.42 | 0.53 | 0.53 | 0.40 ± 0.061 | 66 | 0.70 |
| | mBRAVE | Illumina | $\geq 45$ | 1 | 56.3M | 0.46 | 0.73 | 0.19 | 0.25 ± 0.086 | 86 | 0.86 |
| | MetaWorks | Illumina | $\geq 15$ | 1 | 24.1M | 0.23 | 0.35 | 0.13 | 0.51 ± 0.057 | 50 | 0.58 |
| | QIIME2 | Illumina | $\geq 18$ | 1 | 35.1M | 0.58 | 0.90 | -0.0083 | 0.50 ± 0.045 | 58 | 0.62 |

**Table S4.** Summary biodiversity metrics across filter strategies using the Target + NTS dataset with <u>Genus</u> as the unit of analysis; darker green indicates better performance within each metric and filter strategy.

| Filter Strategy | Software | Dataset | Min. Reads | Min. Reps | # Reads in OTUs | CCC ($\alpha$) | Spearman $\rho$ ($\alpha$) | Mantel $R$ ($\beta$) | BC Dist ± SD ($\beta$) | % Target ($\gamma$) | F1-score ($\gamma$) |
| --- | --- | --- | --- | --- | --- | --- | --- | --- | --- | --- | --- |
| Unfiltered | MAP | Nanopore | 3 | 1 | 52.6M | 0.80 | 0.88 | 0.28 | 0.21 ± 0.12 | 92 | 0.90 |
| | MAP | Illumina | 2 | 1 | 20.3M | 0.18 | 0.62 | 0.13 | 0.47 ± 0.11 | 94 | 0.62 |
| | spcfy.io | Illumina | 1 | 1 | 12.4M | 0.023 | 0.42 | -0.16 | 0.76 ± 0.075 | 56 | 0.37 |
| | mBRAVE | Illumina | 1 | 1 | 20.1M | 0.0017 | 0.28 | -0.14 | 0.79 ± 0.071 | 95 | 0.42 |
| | MetaWorks | Illumina | 3 | 1 | 18.2M | 0.012 | 0.37 | 0.011 | 0.82 ± 0.061 | 51 | 0.32 |
| | QIIME2 | Illumina | 2 | 1 | 14.7M | 0.058 | 0.81 | 0.40 | 0.89 ± 0.033 | 49 | 0.28 |
| Permissive filter ($\geq 0.0001\%$ of sample reads) | MAP | Nanopore | $\geq 3$ | 1 | 52.6M | 0.81 | 0.88 | 0.30 | 0.21 ± 0.12 | 90 | 0.89 |
| | MAP | Illumina | $\geq 2$ | 1 | 20.3M | 0.21 | 0.71 | 0.13 | 0.45 ± 0.11 | 93 | 0.63 |
| | spcfy.io | Illumina | $\geq 1$ | 1 | 12.4M | 0.046 | 0.46 | -0.11 | 0.74 ± 0.083 | 56 | 0.37 |
| | mBRAVE | Illumina | $\geq 1$ | 1 | 20.1M | 0.0028 | 0.12 | -0.13 | 0.75 ± 0.086 | 94 | 0.47 |
| | MetaWorks | Illumina | $\geq 3$ | 1 | 18.2M | 0.011 | 0.30 | 0.012 | 0.82 ± 0.062 | 51 | 0.32 |
| | QIIME2 | Illumina | $\geq 2$ | 1 | 14.7M | 0.053 | 0.73 | 0.25 | 0.89 ± 0.034 | 48 | 0.29 |
| Stringent filter ($\geq 0.001\%$ of sample reads) | MAP | Nanopore | $\geq 12$ | 1 | 52.6M | 0.80 | 0.89 | 0.41 | 0.19 ± 0.11 | 85 | 0.88 |
| | MAP | Illumina | $\geq 10$ | 1 | 20.3M | 0.55 | 0.79 | 0.17 | 0.37 ± 0.11 | 86 | 0.75 |
| | spcfy.io | Illumina | $\geq 5$ | 1 | 12.4M | 0.28 | 0.73 | 0.070 | 0.64 ± 0.084 | 53 | 0.43 |
| | mBRAVE | Illumina | $\geq 10$ | 1 | 20.1M | 0.019 | -0.042 | -0.0023 | 0.60 ± 0.11 | 89 | 0.60 |
| | MetaWorks | Illumina | $\geq 8$ | 1 | 18.2M | 0.048 | 0.41 | -0.14 | 0.77 ± 0.075 | 49 | 0.35 |
| | QIIME2 | Illumina | $\geq 3$ | 1 | 14.7M | 0.063 | 0.26 | 0.30 | 0.89 ± 0.030 | 33 | 0.28 |

**Table S5.** Rankings of pipeline performance on Illumina data (Target + NTS as ground truth). Mean ranks are averaged within diversity categories.

| Filter Strategy | Software | $\alpha$:<br>mean rank | $\beta$:<br>mean rank | $\gamma$:<br>mean rank | Overall Rank |
| --- | --- | --- | --- | --- | --- |
| Unfiltered | MAP | 2.0 | 1.0 | 1.5 | 1 |
|  | spcfy.io | 3.5 | 2.0 | 4.0 | 4 |
|  | mBRAVE | 3.0 | 4.5 | 1.5 | 3 |
|  | MetaWorks | 4.0 | 4.5 | 4.0 | 5 |
|  | QIIME2 | 2.0 | 2.5 | 3.5 | 2 |
| Permissive filter<br>($\geq 0.0001\%$ of sample reads) | MAP | 2.0 | 1.5 | 2.0 | 1 |
|  | spcfy.io | 2.0 | 2.0 | 4.0 | 3 |
|  | mBRAVE | 2.5 | 3.5 | 1.0 | 2 |
|  | MetaWorks | 4.0 | 4.5 | 4.0 | 5 |
|  | QIIME2 | 3.0 | 3.5 | 4.0 | 4 |
| Stringent filter<br>($\geq 0.001\%$ of sample reads) | MAP | 2.5 | 3.5 | 2.0 | 2 |
|  | spcfy.io | 3.5 | 2.0 | 4.0 | 4 |
|  | mBRAVE | 1.5 | 1.5 | 1.0 | 1 |
|  | MetaWorks | 3.0 | 4.0 | 4.0 | 5 |
|  | QIIME2 | 2.5 | 3.5 | 3.0 | 3 |

**Table S6.**
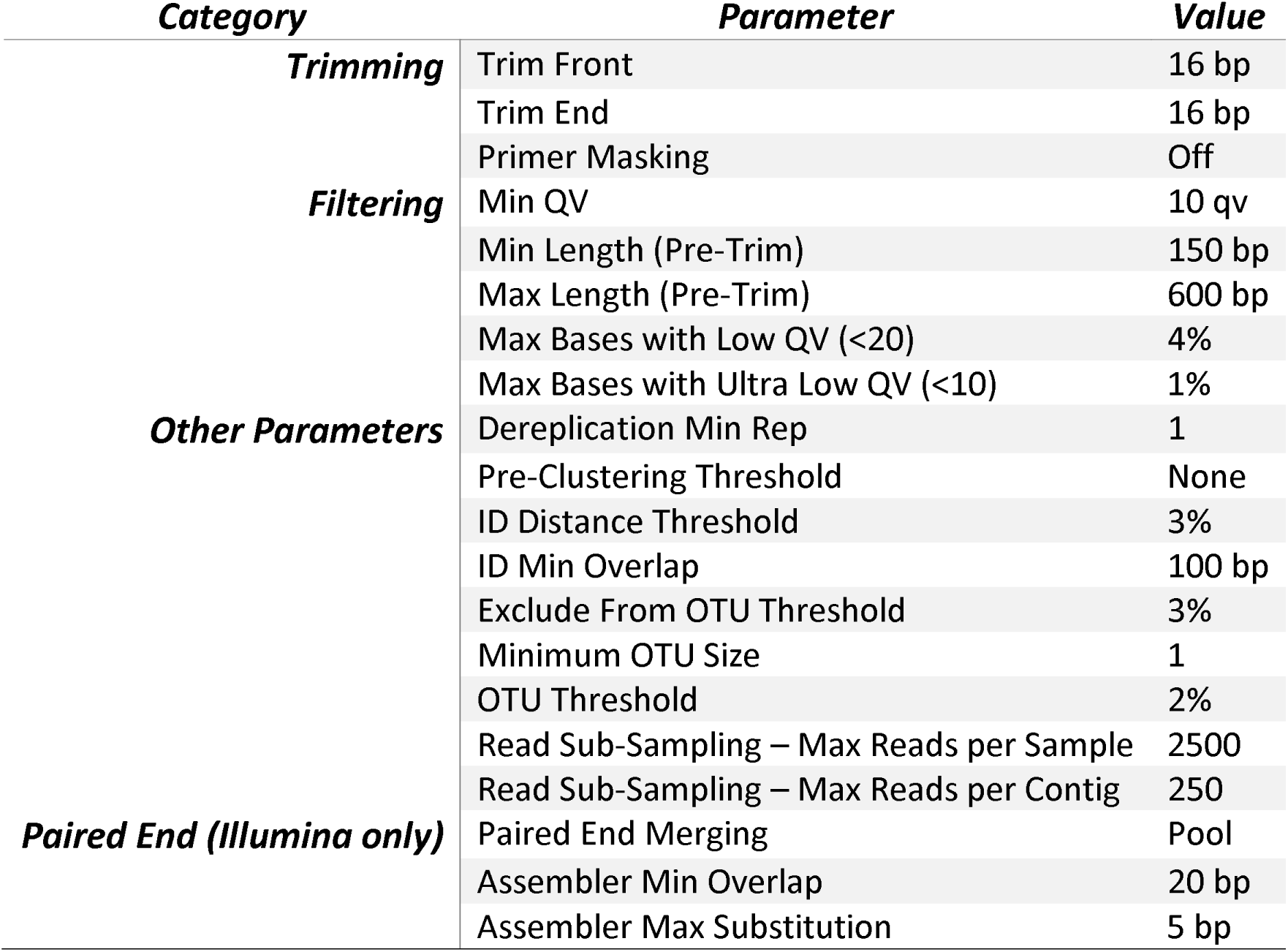
mBRAVE project parameters.

